# The impact of exercise intensity on neurophysiological indices of food-related inhibitory control and cognitive control: A randomized crossover event-related potential (ERP) study

**DOI:** 10.1101/2020.11.10.377267

**Authors:** Bruce W. Bailey, Alexandra M. Muir, Ciera L. Bartholomew, William F. Christensen, Kaylie A. Carbine, Harrison Marsh, Hunter LaCouture, Chance McCutcheon, Michael J. Larson

## Abstract

Food-related inhibitory control, the ability to withhold a dominant response towards highly palatable foods, influences dietary decisions. Food-related inhibitory control abilities may increase following a bout of aerobic exercise; however, the impact of exercise intensity on both food-related inhibitory control and broader cognitive control processes is currently unclear. We used a high-powered, within-subjects, crossover design to test how relative intensity of aerobic exercise influenced behavioral (response time, accuracy) and neural (N2 and P3 components of the scalp-recorded event-related potential [ERP]) measures of food-related inhibitory and cognitive control. Two hundred and thirteen participants completed three separate conditions separated by approximately one week in randomized order: two exercise conditions (35% [moderate] or 70% [vigorous] of VO_2max_) and seated rest. Directly following exercise or rest, participants completed a food-based go/no-go task and a flanker task while electroencephalogram data were recorded. Linear mixed models showed generally faster response times (RT) and improved accuracy following vigorous exercise compared to rest, but not moderate-intensity exercise; RTs and accuracy did not differ between moderate intensity exercise and rest conditions. N2 and P3 amplitudes were larger following vigorous exercise for the food-based go/no-go task compared to rest and moderate intensity exercise. There were no differences between exercise conditions for N2 amplitude during the flanker task; however, P3 amplitude was more positive following vigorous compared to rest, but not moderate exercise. Gender did not moderate exercise outcomes. Results suggest improved and more efficient food-related recruitment of later inhibitory control and cognitive control processes following vigorous exercise.

## 1. Introduction

Cognitive control is the ability to allocate neural resources in order to adapt and interact with the surrounding environment and update behavior to achieve one’s goals (Mackie et al., 2013; Miller and Cohen, 2001). Cognitive control encompasses a variety of component processes, including performance monitoring, inhibitory control, and attention/control allocation, all which involve interplay between the anterior cingulate and prefrontal cortices, among other areas (Botvinick et al., 2001; Miller and Cohen, 2001). Although cognitive control is essential for goal-directed behaviors, the factors that enhance or decrease cognitive control abilities are areas of continued research.

One factor that may impact an individual’s cognitive control abilities is exercise, with recent literature demonstrating that exercise may have a small enhancing effect on cognitive control component processes (for example Chang et al., 2012; Guiney and Machado, 2013; Kao et al., 2019; Kempton et al., 2011; Ligeza et al., 2018; Ludyga et al., 2016). This general improvement in cognitive control abilities following exercise suggests that, in addition to improving overall physical and mental health (Knapen et al., 2015; LeBouthillier and Asmundson, 2017; Morres et al., 2019), exercise may also acutely improve our ability to accurately identify environmental demands to achieve goal-directed behavior.

Although there appears to be a small positive effect of exercise on general cognitive control abilities, not all results are in agreement (Lambourne and Tomporowski, 2010; Tomporowski and Ellis, 1986). This heterogeneity in findings may be partially due to the variety of exercise intensities employed in studies, as different intensities of exercise could have different effects on subsequent cognitive control (Olson et al., 2016; Wohlwend et al., 2017). Meta-analytic evidence shows that light-to-moderate intensity exercise has a small, but beneficial, effect on cognitive control; however, this positive effect is only present immediately following the acute exercise (Chang et al., 2012). In comparison, moderate-to-vigorous exercise demonstrates the same small positive effect on cognitive control, but the effect lasts longer when compared to lighter intensity exercise (Chang et al., 2012; Moreau and Chou, 2019). Taken together, although exercise may have a small positive effect on cognitive control, the intensity at which exercise is performed may differentially affect subsequent cognitive control abilities and the length to which the effect extends.

One facet of cognitive control that may be particularly influenced by exercise intensity is inhibitory control. Inhibitory control is the ability to withhold a dominant response to override basic instincts or habits to produce goal directed behavior (Diamond, 2013; Ko and Miller, 2013). A single bout of aerobic exercise acutely enhances inhibitory control abilities (Kamijo et al., 2007; Kao et al., 2017). This enhancement of inhibitory control abilities may be attributed to increased blood flow in the dorsolateral prefrontal cortex during or directly following exercise (Byun et al., 2014; Yanagisawa et al., 2010). There is also evidence that exercise may increase general oscillatory brain activity when compared to a resting state, causing an enhancement of multiple cognitive processes rather than inhibitory control abilities specifically (Ciria et al., 2018). Further research surrounding the role of exercise intensity in inhibitory control abilities is needed to parse apart what exact intensities of exercise may enhance inhibitory control (Carbine, In Press).

Event-related potentials (ERP) derived from electroencephalogram (EEG) data can be utilized to understand the neural bases of cognitive and inhibitory control, including how acute bouts of aerobic exercise affect these cognitive processes. One neural index of inhibitory control is the N2 component of the scalp-recorded ERP. The N2 is a negative-going ERP that peaks approximately 200 to 350 milliseconds following the onset of a stimulus. N2 amplitude becomes more negative as additional inhibitory resources are recruited to withhold a dominant response towards a stimulus (Folstein and Van Petten, 2008; Larson et al., 2014). The cognitive process which the N2 indexes depends on the task and stimuli being utilized for the experiment at hand, with the N2 reflecting response inhibition during both a go/no-go (Folstein and Van Petten, 2008) and flanker tasks (Van Veen and Carter, 2002; Xie et al., 2017). In addition to the N2, the P3 is a positive-going waveform that appears approximately 300 to 600 ms following the presentation of a stimuli, whether auditory or visual (Falkenstein et al., 1999). The P3 is larger when inhibiting a dominant motor response (Gajewski and Falkenstein, 2013) and when suppressing attention towards other nonrelevant stimuli in the environment (Polich, 2007). The functional significance of the P3 is still being debated, however, prominent theories posit that the P3 component is representative of context updating following stimuli or is representative of the allocation of attentional resources to salient stimuli (Polich, 2007).

The few studies that have examined the impact of exercise intensity on inhibitory control processes reflected by the N2 ERP component report mixed results. Larger (more negative) N2 amplitudes were observed during exercise (at 40% and 60% of VO_2_ peak) compared to a seated rest condition, suggesting greater cognitive control implementation during exercise (Olson et al., 2016). However, the effects of exercise on N2 amplitude may be different when measured directly following exercise with N2 amplitude decreasing following moderate exercise (60% of max heart rate) in both adults and children (Pontifex and Hillman, 2007; Stroth et al., 2009). Ligeza et al. (2018) observed differential effects of exercise intensity in a between-subjects design, with N2 amplitude becoming larger at submaximal aerobic intensity (between the first and second ventilatory thresholds) when compared to rest, but smaller after high intensity interval training compared to rest (Ligeza et al., 2018). Taken together, these results suggest that exercise intensity may play a role in inhibitory control as indexed by N2 amplitude, although the direction of that relationship is currently unclear.

Although the findings surrounding the inhibitory control processes reflected by the N2 and exercise are variable, the variability in quantifying and implementing intensity of exercise may at least partially explain the heterogeneity in results. Themanson et al. (2006) had participants exercise at 85% of their maximal heart rate, which is considered to be high intensity exercise. Pontifex and Hillman (2007) along with Stroth et al. (2009) used 60% of the individual’s estimated maximum heart rate to define moderate exercise, while Ligeza et al., (2018) used ventilatory thresholds. As Ligeza et al. explains, this wide variety in definitions for exercise intensity may cause each study to be examining different intensities of exercise per participant. As such, standardized methods of exercise intensity based on the physical fitness of the individual participant is essential to understand how exercise intensity affects cognition.

Similar to the N2, acute exercise seems to have mixed effects on cognitive control processes reflected by P3 amplitude. In the most exhaustive meta-analysis to date, Kao et al. (2020) concluded that P3 amplitude generally increases following an acute bout of continuous aerobic submaximal exercise when compared to rest. However, for some studies, P3 amplitude after aerobic exercise was moderated by the age of the participant (Brush et al., 2020; Kamijo and Takeda, 2009; Lennox et al., 2019), fitness level (Tsai et al., 2016; Tsai et al., 2014), emotional context of exercise (Miller et al., In Press), and baseline levels of inhibitory control abilities (Drollette et al., 2014), suggesting multiple moderating factors in the relationship between exercise and P3 amplitude (see also Chacko et al., 2020). Interestingly, across various studies included in the meta-analysis, larger P3 amplitude was observed when comparing continuous aerobic exercise to high intensity interval training and rest, while decreased P3 amplitude was observed between high intensity interval training and rest (Kao et al., 2017). As outlined by Kao et al. (2020), in general, it seems as if continuous aerobic exercise may be beneficial to inhibitory control processes (Hillman et al., 2003; Kamijo et al., 2007; O’Leary et al., 2011), but much like the N2, this relationship may be different depending on the intensity of exercise.

Moderation of dietary behavior is one specific example of the importance of inhibitory control. Despite widespread evidence that a healthy diet reduces the risk of obesity, Type 2 diabetes, cardiovascular disease, high blood pressure, and depression to name a few (Carek et al., 2011; Cornelissen and Smart, 2013; Fiuza-Luces et al., 2018; Kirwan et al., 2017; Swift et al., 2018), the impulse to consume highly palatable and high-calorie foods is difficult to inhibit, even if an individual has recently eaten (Armelagos, 2014; Rogers and Brunstrom, 2016). This is complicated in the current environment where highly palatable, high calorie foods are plentiful and food related cues (food related pictures, ads and smells) are ubiquitous. As such, highly palatable and high-calorie foods require specific inhibitory control to reduce automatic urges to consume (Carbine et al., 2017; Guerrieri et al., 2007).

Individuals who are obese may display lower inhibitory control (Lavagnino et al., 2016; Spitoni et al., 2017), suggesting a decreased ability to withhold the dominant response to moderate caloric intake. In addition, lower inhibitory control is associated with overeating (Guerrieri et al., 2007), along with higher consumption of carbohydrates and more calories overall (Ko and Miller, 2013). Inhibitory control predicts saturated fat intake, rather than the consumption of fruits and vegetables, suggesting that inhibitory control is involved in the withholding of dietary behavior rather than the initiation of eating healthy foods (Allom and Mullan, 2014).

As both the N2 and P3 ERP components can be used to index general inhibitory control, both of these event-related potentials can also be used to index food-related inhibitory control. The N2 is more negative as an individual inhibits a response to food stimuli when compared to non-food stimuli (Watson and Garvey, 2013) and more negative when inhibiting to high-calorie foods when compared to low-calorie foods (Carbine et al., 2017; Carbine et al., 2018b). These results suggest an increased need for inhibitory control neural resources when inhibiting a response towards high-calorie foods. Similar to the N2, P3 amplitude becomes larger (i.e., more positive) when inhibiting towards high-calorie foods compared to low-calorie foods (see also Aulbach et al., 2020; Carbine et al., 2017; Carbine et al., 2018a), again suggesting increased cognitive control when inhibiting to high-calorie foods.

As a number of studies have demonstrated a relationship between exercise intensity and inhibitory control, it is possible that exercise intensity also moderates the relationship between exercise and food-specific inhibitory control. Generally, researchers have hypothesized that physical activity may indirectly affect eating behavior through strengthening the neural circuits in the prefrontal cortex that influence inhibitory control, which in turn reduces impulses to consume high-calorie foods (Joseph et al., 2011). In one study, after a bout of aerobic exercise, inhibitory control increased (as indexed by accuracy and response time), and subsequently reduced consumption of high-calorie foods directly following the completion of the exercise condition (Lowe et al., 2016). In addition, several studies have observed reduced food intake acutely following exercise (Hagobian et al., 2013; Schubert et al., 2013; Sim et al., 2014). However, a gap in the literature is research that has rigorously tested the neural mechanisms of food-related inhibitory control following different levels of exercise intensity.

### 1.1 Aims and hypotheses

Previous studies examining the relationship between exercise and inhibitory control have generally focused on how one intensity of exercise differs from rest, rather than examining how different intensities of exercise differentially effect cognitive control in the same sample of participants. Additionally, although there have been a number of studies that have examined the relationship between exercise and cognitive control, how this relationship extends to food-specific inhibitory control is less known. As such, the current study used a within-subjects crossover, design to evaluate the impact of moderate and vigorous exercise on both cognitive control and food-related inhibitory control. Given blood-flow based neuroimaging studies that suggest increased cerebral blood flow perfusion during mild-to-moderate intensity exercise, with a subsequent decrease toward resting values during vigorous exercise likely because of vasoconstriction during high intensity exercise (Joris et al., 2018; Ogoh and Ainslie, 2009), we hypothesized that there would be an inverted U-shaped relationship between both food specific inhibitory control and general inhibitory control. Specifically, we hypothesized that N2 and P3 amplitude would increase for moderate intensity exercise but decrease for high intensity exercise when compared to seated rest for both the food-specific and general cognitive control tasks.

## 2. Materials and Method

All data for the current study are available on a study-specific Open Science Framework webpage: https://osf.io/u9bdy/.

### 2.1 Participants

All experimental procedures were approved by the Institutional Review Board at Brigham Young University and participants provided written informed consent. Exclusion criteria were determined by participant self-report and included being diagnosed with an eating disorder, psychiatric disorder, head injury resulting in loss of consciousness, a body mass index below 18.5, current pregnancy or lactation, or more than 225 minutes of cardiorespiratory exercise on average per week. Participants were between 18 and 45 years of age and self-endorsed the ability to exercise at a vigorous intensity (i.e., jog for 40 minutes). Prior to study enrollment, the Physical Activity Readiness Questionnaire (PAR-Q; (Arraiz et al., 1992) was used to screen the participant’s ability to participate in physical activity. If any item was endorsed on the readiness questionnaire then the participant was not enrolled.

The sample size for the current study was calculated *a priori* (see Larson and Carbine, 2017) based on a previous study examining the effects of exercise on attention to visual food cues in obese and normal weight women. In the Carbine et al. study, we observed a mean standard error of 2.77 µV between exercise intensity conditions which was used in the current power analysis. With alpha set at .05 and beta at .80, we would need a sample of 200 participants to detect a mean difference as small as .43 µV, which represents a 25% difference between the rest and 70% vigorous-intensity exercise conditions. Thus, we recruited 230 participants for the current study due to an *a priori* estimated 15% dropout rate.

Participant characteristics are described in Table 1. A total of 462 men and women were assessed for eligibility and 230 were randomized to exercise condition order (see Figure 1). Of the 230 participants that were randomized, 212 finished the study. Participants who did not complete the study cited loss of interest and lack of time to commit to the study as reasons for discontinuing participation. Those who did not finish the study did not differ from those who finished the study on key demographic characteristics that included age (*t*(24.34) = −0.11, p = 0.91), body mass index (*t*(20.24) = −1.71, p = 0.10), body fat (*t*(19.97) = −0.21, p = 0.83) or VO_2max_ (t(18.78) = −0.42, p = 0.68). Heart rate and metabolic equivalent exercise intervention characteristics for the two exercise conditions are presented in Table 2.

**Table 1:** Mean and standard deviation of height, weight and VO_2max_ (n = 217)

|  |  | Mean | SD | Range |
| --- | --- | --- | --- | --- |
| **Age (years)** | **Total** | **22.46** | **3.68** | **18, 44** |
|  | Men | 23.14 | 3.65 | 18, 44 |
|  | Women | 21.78 | 3.58 | 18, 39 |
| **Height (cm)** | **Total** | **171.73** | **9.46** | **148, 195** |
|  | Men | 178.31 | 6.77 | 161.9, 195 |
|  | Women | 164.77 | 6.43 | 148, 179.50 |
| **Weight (kg)** | **Total** | **69.22** | **14.07** | **45.30, 129.30** |
|  | Men | 76.15 | 13.73 | 57.10, 129.30 |
|  | Women | 61.89 | 10.22 | 45.30, 93.35 |
| **VO<sub>2max</sub> (ml kg<sup>-1</sup> min<sup>-1</sup>)** | **Total** | **44.05** | **7.80** | **28.21, 67.87** |
|  | Men | 48.36 | 7.42 | 28.21, 67.87 |
|  | Women | 39.50 | 5.17 | 28.4, 52.95 |
| **BMI (kg m<sup>-2</sup>)** | **Total** | **23.29** | **3.24** | **17.60, 36.90** |
|  | Men | 23.80 | 3.44 | 17.60, 36.90 |
|  | Women | 22.74 | 2.93 | 18.00, 33.50 |
| **Max Heart Rate** | **Total** | **195** | **9.01** | **169, 225** |
|  | Men | 193 | 9.28 | 169, 225 |
|  | Women | 196 | 8.60 | 175, 214 |
| **Body fat (%)** | **Total** | **25.97** | **8.40** | **9.69, 48.74** |
|  | Men | 20.73 | 6.92 | 17.37, 48.85 |
|  | Women | 31.51 | 5.91 | 9.70, 45.18 |

SD = standard deviation.

**Figure 1:**
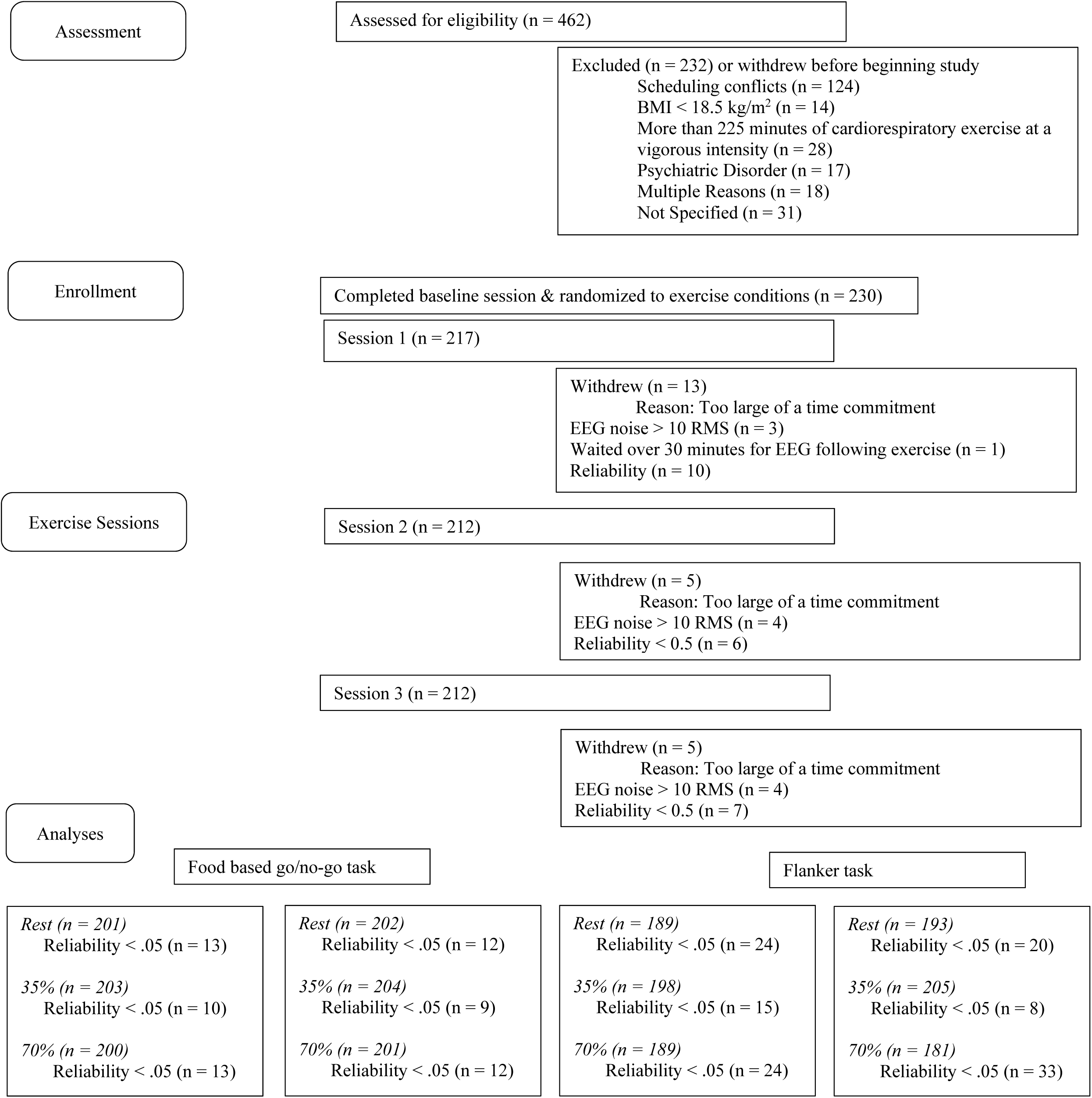
Overview of participant recruitment and analyses. RMS = root mean squared. As a note, numbers presented as being dropped for reliability are re-presented under analyses to show the distribution of data loss across exercise condition rather than session.

**Table 2:** Heart rate and metabolic equivalent characteristics of exercise by condition

|  | 35% |  | 70% |  |
| --- | --- | --- | --- | --- |
|  | Mean | SD | Mean | SD |
| Average Heart Rate |  |  |  |  |
| Male | 107.9 | 12.2 | 162.2 | 11.4 |
| Female | 110.2 | 12.9 | 163.2 | 11.4 |
| % of Max Heart Rate |  |  |  |  |
| Male | 56.7% | 5.6% | 82.9% | 5.0% |
| Female | 55.1% | 5.9% | 83.9% | 5.7% |
| METs |  |  |  |  |
| Male | 55.1 | 5.4 | 82.9 | 4.7 |
| Female | 56.9 | 5.9 | 84.4 | 4.8 |

MET = Metabolic equivalent; 1 MET = resting metabolism
< 3 METs = light intensity activity, 3 to 6 METs = moderate intensity exercise,
> 6 METs = vigorous intensity exercise. SD = standard deviation.

### 2.2 Procedures overview

Each participant completed four lab visits, which included baseline testing in addition to three separate conditions conducted in randomized order: vigorous-intensity exercise (70% of VO_2max_), moderate-intensity exercise (35% of VO_2max_), and seated rest. At the baseline session, participants were measured for height, weight, and body composition. They then completed a volitional fatigue VO_2max_ test that was used to prescribe the exercise interventions for the future 35% and 70% VO_2max_ conditions. Baseline measurements took place at least two days prior to completion of subsequent study conditions.

The order of the three experimental sessions (70% VO_2max_, 35% VO_2max_, rest) were randomly assigned using a random number generator. Prior to arrival for each session, participants endorsed that they adhered to each of the following: slept for at least seven hours the night before, were adequately hydrated, did not consume any food or beverages with caloric content in the four hours preceding the session, and did not consume caffeine or perform vigorous exercise during the 24 hours preceding the session (see Carbine et al., 2017). Each of the three conditions were administered the same day of the week, at the same time of day, one week apart. If a participant could not attend on their scheduled day they were re-scheduled for the following week to maintain day of week and time of day consistency.

During the rest condition, instead of exercising participants were seated while they completed a battery of questionnaires that included the Dutch Eating Behavior Questionnaire, the Yale Food Addiction Scale, and the Depression Anxiety Stress Scale. The data from these questionnaires were collected but are not reported in the current study as they were not part of the current hypotheses and are only included here for transparency/completeness of presentation. After completing the battery of questionnaires, the participants watched a 40-minute documentary titled “What Plants Talk About” (https://www.youtube.com/watch?v=cIftMUWs4q0) so they were in the lab a similar amount of time prior to the EEG recording as the exercise sessions, but in a resting situation watching a low-arousal video.

After the completion of the exercise or rest bout, participants were escorted up to the electroencephalogram research suite. The participant was given a towel to dry off while a fan blew to reduce sweat. Then, the participant was fit with an EEG cap, after which they completed a food-based go/no-go task and a flanker task (always in that order) during which EEG data were recorded. Upon completion of the computerized tasks, the EEG cap was removed and participants were either given a calendar reminder for their next session date or were compensated $40 or provided course credit at the completion of the study.

### 2.3 Measurements

#### 2.3.1 Anthropometrics

For descriptive purposes, height, weight and body composition were measured. Height was measured to the nearest 0.1 cm using a wall mounted stadiometer (SECA, Chino, California) and weight was measured to the nearest 0.1 kg (Tanita, Arlington Heights, IL). GE iDXA was used to describe body composition (GE, Fairfield, CT).

#### 2.3.2 VO_2max_ Test

VO_2max_ was determined using a modified George protocol (George, 1996). Safety guidelines outlined by the American College of Sports Medicine were followed to ensure participant safety (ACSM, 2018). Participants started the test with a 7-minute warm-up ona treadmill walking at 3 mph with a grade that increased from 0-6%. Then the grade was lowered to 0% and participants chose a comfortable running speed at which to complete the rest of the test. Perceived exertion was measured using the Borg 6-20 scale after every minute of exercise and heart rate was measured continuously throughout the testing using a FT7 Polar heart rate monitor (NY, USA). After three minutes running at the selected pace, the grade increased by 1.5% every minute. The test stopped when the participant self-reported voluntary exhaustion. The test was considered maximal if three of the five following criteria were met: the participant physically could not continue, their perceived exertion was either 19 or 20 on the Borg scale, their heart rate was within 15 beats per minute of their predicted maximum, their VO_2_ began to plateau, or their respiratory exchange ratio was ≥ 1.0. Participants concluded the test with a four-minute walking cool-down. Measurements were taking using the COSMED Quark Ergo metabolic cart (Chicago, IL).

During the moderate-intensity and vigorous exercise sessions, participants jogged at a specified percentage (70% or 35%) of their VO_2max_ calculated at their baseline session. The intensity was prescribed directly based on the participant’s measured maximal capacity. Within the first five minutes of the session, participants gradually increased exercise intensity until their specified percentage of VO_2max_ was achieved. They then maintained the intensity for the remainder of the exercise session (35 minutes). The exercise bout lasted for 40 minutes, including the buildup to the specified percentage of VO_2max_. As such, all three pre-EEG activities (rest, moderate, and vigorous) lasted 40 minutes prior to EEG net application and completing the computerized tasks. If the participant needed to stop and take a break at any point during the exercise bout, the time was paused and continued after the participant began exercising again.

### 2.4 Computerized tasks

#### 2.4.1 High-calorie go/no-go tasks

Participants were instructed to respond with a button press when they saw a low-calorie food (go-trial) and withhold all responses when a high-calorie food was presented (no-go trial). All stimuli were presented in a random order. Participants completed two blocks of 100 trials each, with 70 go trials and 30 no-go trials. This distribution of go/no-go trials was used to establish a predominance of go trials, making inhibitory behavior more challenging. Pictures of low and high-calorie foods were separated by a fixation cross jittered randomly from 600 to 700 milliseconds. Stimuli remained on the screen for 500 ms, and responses made after 1000 ms were considered omission errors and not used in data analyses.

Pictures used for the food stimuli were provided by Killgore and colleagues (2003) who have used these same images in papers published previously (e.g., Killgore et al., 2013; Killgore and Yurgelun-Todd, 2005, 2007). These images were first categorized by 26 separate undergraduates who rated all 120 pictures as either high- or low-calorie foods. Only stimuli that were accurately categorized as high- and low-calorie foods at least 95% of the time were used, resulting in 38 pictures for each category (see Carbine et al., 2017). Low-calorie food stimuli included 13 vegetables and 25 fruits. High-calorie food images consisted of 16 desserts, 15 high-calorie dinner meals, and 7 high-calorie breakfast meals. This task has been used previously and consistently elicits a more negative N2 and more positive P3 towards high-calorie foods (no-go trials) when compared to low-calorie foods (go trials; Carbine et al., 2017; Carbine et al., 2018a).

#### 2.4.2 Flanker task

Upon completion of the go/no-go task, participants completed a modified arrow version of the Ericksen flanker task (Eriksen and Eriksen, 1974). Participants were instructed to respond as quickly and accurately as possible by pressing a button that corresponded to the directionality of the middle arrow. Congruent (e.g., < < < < <) and incongruent (e.g., < < > < <) arrow groups in 36-point Arial white font were randomly presented in the center of a black screen. To establish pre-potency, flanking arrows were presented for 100 ms prior to the onset of the middle arrow, which remained on the screen for an additional 600 ms. If a participant responded after 1,000 ms, the response was considered an error of omission and was not included in analyses. Between each trial, a fixation cross was shown for either 800 ms, 1,000 ms, or 1,200 ms. These three fixation cross intervals were split evenly across the 204 trials. Two blocks of 102 trials each were completed with 44% of trials being congruent and 56% of trials being incongruent.

### 2.5 EEG data acquisition and reduction

EEG data were collected and are reported according to the guidelines for studies using electroencephalography (Clayson et al., 2019; Keil et al., 2014). Specifically, all EEG data were collected from 128 equidistant passive Ag/AgCl electrodes in a hydrocel geodesic sensor net using an Electrical Geodesics, Inc. series 300 amplifier (20K nominal gain, band-pass = 0.01-100 Hz). All data were referenced to the vertex electrode during data collection and were digitized continuously at 250 Hz with a 16-bit analog to digital converter. Electrode impedances were kept at or below 50 kΩ per the manufacture’s recommendation. Offline, following data collection, data were digitally filtered with a 0.1 Hz high pass filter (0.3 rolloff; 36.9 db/octave) and 30 Hz low pass filter (0.3 rolloff; 19.5 db/octave) in NetStation (v5.3.0.1). Data were subsequently epoched from 200 ms before stimulus onset to 1000 ms following stimulus onset for both the flanker and the food-based go/no-go tasks. For the go/no-go task, trials were segmented to include only correct go and no-go trials. For the flanker task, trials were segmented to include only correct congruent and incongruent trials. Eye movements and blink artifacts were then corrected using independent components analysis (ICA) in the ERP PCA toolkit (Dien, 2010). If a component correlated with two blink templates (one from the ERP PCA toolkit and the other derived by the authors) at a level of 0.9 or higher, that component was subsequently removed from the data. If any electrode had a fast average amplitude of over 50 microvolts or if the fast average amplitude was greater than 100 microvolts, the channel was defined as bad and replaced using the nearest six electrodes for interpolation (Dien, 2010).

Following artifact correction, data were average re-referenced and baseline adjusted from 200 ms before stimulus onset using the ERP PCA toolkit (Dien, 2010). For the food-based go/no-go task, data were analyzed from a region of interest in the frontocentral area consisting of four a priori chosen electrodes (electrodes 6 [FCz], 7, 106, and 129 [Cz]; Carbine et al. (2017); Carbine et al. (2018a); see Larson, Farrer, & Clayson, (2011a) for electrode montage). Time windows were determined using a collapsed localizer approach over the region of interest wherein we visually examined the grand average waveforms collapsed across all conditions to determine the appropriate time window (Luck and Gaspelin, 2017). Mean amplitude for the N2 was extracted from 200 to 300 ms following stimulus onset, while P3 mean amplitude was extracted from 400 to 550 ms following stimulus onset (see Carbine et al. (2017) and Carbine et al. (2018a) for similar time windows).

For the flanker task, N2 amplitude was analyzed from the same *a priori* chosen frontocentral region of interest (electrodes 6, 7, 106, and 129) and P3 amplitude was analyzed from a frontomedial region of interest consisting of four *a priori* selected electrodes (electrodes 129, 31, 55, 80 (Larson et al., 2011a)). A collapsed localizer approach (collapsing across all conditions) over each region of interest was again used to select time windows. The N2 amplitude was extracted using an adaptive mean amplitude of 16 ms from 270 ms to 380 ms following target arrow onset while the P3 amplitude was extracted using a mean amplitude from 370 to 500 ms following target arrow onset. Mean amplitude was used along with region of interests due to evidence suggesting that averaging multiple electrodes together increases signal reliability when compared to a single electrode (Clayson, 2020; Clayson et al., 2013).

### 2.6 Reliability analysis

To determine the minimum number of trials necessary to achieve adequate reliability for the N2 and P3 components, dependability estimates of ERPs were assessed through the ERP Reliability Analysis Toolbox v.0.3.2 (Clayson and Miller, 2017) using generalizability theory. To meet assumptions of independent colinearity, dependability estimates were calculated and are reported separately for each condition. Minimum dependability cut-offs were set at 0.5 (although overall dependability ranged from 0.64 to 0.96), and therefore, any participant that did not meet the dependability cut-off of 0.5 was taken out from further data analyses. For specific dependability estimates and minimum and maximum trial numbers by condition and ERP component, see Table 3.

**Table 3:** Dependability estimates and trial ranges for all ERP components

|  | Rest |  | 35% |  | 70% |  |
| --- | --- | --- | --- | --- | --- | --- |
|  | Estimate | Trial Range | Estimate | Trial Range | Estimate | Trial Range |
| Flanker N2 |  |  |  |  |  |  |
| Congruent | 0.83 | 49, 120 | 0.77 | 34,120 | 0.73 | 38,120 |
| Incongruent | 0.72 | 38,114 | 0.71 | 41, 113 | 0.78 | 31, 113 |
| Flanker P3 |  |  |  |  |  |  |
| Congruent | 0.84 | 29, 120 | 0.82 | 28, 120 | 0.64 | 59, 120 |
| Incongruent | 0.84 | 22, 114 | 0.82 | 22, 113 | 0.84 | 46, 113 |
| GNG N2 |  |  |  |  |  |  |
| Go | 0.96 | 12, 140 | 0.96 | 16, 140 | 0.95 | 14, 140 |
| No-go | 0.88 | 8, 59 | 0.93 | 5, 58 | 0.90 | 8, 60 |
| GNG P3 |  |  |  |  |  |  |
| Go | 0.94 | 12, 140 | 0.92 | 31, 140 | 0.92 | 13, 140 |
| No-go | 0.91 | 5, 59 | 0.89 | 8, 58 | 0.90 | 8, 60 |

For the go/no-go task, 36 sessions were removed for the N2 component (5.2% of all sessions, 13 [75% condition], 10 [35% condition], 13 [rest condition]) while 33 sessions were removed for the P3 component (4.8% of all sessions, 12 [70% condition], 9 [35% condition], 12 [rest condition]). Fifty additional sessions were removed due to participant not completing a session or computer malfunction. Thus, the final sample size for the N2 was 200 sessions for 70% exercise, 203 sessions for 35% exercise, and 201 sessions for rest. The final number of sessions for the P3 included 201 sessions for 70% exercise, 204 sessions for 35% exercise, and 202 sessions for rest. Overall, dependability estimates for each condition were above 0.71 for the N2 and above .64 for the P3, suggesting adequate reliability for both ERP components.

For the N2 component derived from the flanker task 63 sessions were removed for not meeting the minimum 0.5 reliability threshold (9.1% of all sessions, 24 [70% condition], 15 [35% condition], 24 [rest condition]) while 61 sessions were removed for the P3 component (8.8% of all sessions, 33 [70% condition], 8 [35% condition], 20 [rest condition]). Additionally, 51 sessions were excluded from data analyses due to the participant not completing a session or computer malfunction. Thus, for the N2, 189 70% exercise sessions, 198 35% exercise sessions, and 189 rest sessions were included in the final analyses. The P3 analyses included 181 70% exercise condition sessions, 205 35% condition sessions, and 193 rest condition sessions.

### 2.7 Behavioral data

Mean accuracy and median response time were extracted for both the food-based go/no-go task and the flanker task. Both mean accuracy and median response time (RT) were separated as a function of trial-type (go/no-go, congruent/incongruent) and exercise/rest condition. Mean accuracy and median RT separated by exercise condition are presented in Tables 4 and 5.

**Table 4:** Behavioral and ERP means and standard errors for the food-related go/no-go task by condition

|  | Rest |  | 35% |  | 70% |  | F | p |
| --- | --- | --- | --- | --- | --- | --- | --- | --- |
|  | Mean | SE | Mean | SE | Mean | SE |  |  |
| Response Time (ms) |  |  |  |  |  |  | 6.52 | 0.002* |
| Go | 4.35 | 2.83 | 5.82 | 2.83 | 10.17 | 2.83 |  |  |
| Response Accuracy (%) |  |  |  |  |  |  | 1.13 | 0.32 |
| Go | 97.42 | 0.52 | 98.23 | 0.52 | 98.64 | 0.52 |  |  |
| No-go | 87.78 | 0.52 | 88.64 | 0.52 | 89.58 | 0.52 |  |  |
| N2 Mean Amplitude (μV) | | | | | | | 0.09 | 0.92 |
| Go | -1.46 | 0.15 | -1.39 | 0.15 | -1.79 | 0.15 |  |  |
| No-go | -1.98 | 0.15 | -1.95 | 0.15 | -2.35 | 0.15 |  |  |
| P3 Mean Amplitude (μV) | | | | | | | 3.56 | 0.03* |
| Go | 1.19 | 0.14 | 1.21 | 0.14 | 1.34 | 0.14 |  |  |
| No-go | 2.40 | 0.14 | 2.40 | 0.14 | 2.90 | 0.14 |  |  |

Note: Values presented for F and p refer to the exercise condition-by-trial type interaction except for the response time, where it is just the main effect of condition
\*p < .05, SE = standard error

**Table 5:** Behavioral and ERP means and standard errors for the flanker task by condition

|  | Rest |  | 35% |  | 70% |  | F | p |
| --- | --- | --- | --- | --- | --- | --- | --- | --- |
|  | Mean | SE | Mean | SE | Mean | SE |  |  |
| Response Time (ms) |  |  |  |  |  |  | 5.28 | 0.005* |
| Congruent | 353.40 | 2.31 | 355.75 | 2.32 | 351.59 | 2.32 |  |  |
| Incongruent | 419.11 | 2.31 | 419.11 | 2.32 | 411.03 | 2.32 |  |  |
| Response Accuracy (%) |  |  |  |  |  |  | 1.13 | 0.33 |
| Congruent | 96.04 | 0.52 | 95.02 | 0.53 | 94.78 | 0.53 |  |  |
| Incongruent | 88.71 | 0.52 | 87.92 | 0.53 | 88.42 | 0.53 |  |  |
| N2 Mean Amplitude (μV) | | | | | | | 0.82 | 0.44 |
| Congruent | 1.27 | 0.14 | 1.38 | 0.13 | 1.41 | 0.14 |  |  |
| Incongruent | 0.24 | 0.14 | 0.16 | 0.13 | 0.20 | 0.14 |  |  |
| P3 Mean Amplitude (μV) | | | | | | | 0.30 | 0.74 |
| Congruent | 2.70 | 0.16 | 3.00 | 0.16 | 3.02 | 0.17 |  |  |
| Incongruent | 4.05 | 0.16 | 4.30 | 0.16 | 4.49 | 0.17 |  |  |

Note: Values presented for F and p refer to the exercise condition by trial-type interaction
\*p < .05, SE = standard error

### 2.8 Statistical analyses

Means and standard errors are reported for all variables of interest in Tables 4 and 5. Alpha for statistical tests was set at 0.05. To determine how exercise intensity affected both behavioral measures (accuracy and RT) and neural measures (N2 amplitude and P3 amplitude) of inhibitory control, eight separate linear mixed models were fit in PC-SAS (v. 9.4). Condition (seated rest, 35% of VO_2max_, and 70% of VO_2max_) and trial-type (go vs. no-go or congruent vs. incongruent) were the fixed effects and participant the random effect. The interaction between condition and trial-type was evaluated for all models, except the model evaluating go/no-go response time. This model only evaluated the main effect of condition since there was no response time associated with the no-go (withhold response) trials. The LSmeans procedure was used to evaluate significant main and interactive affects. The Tukey-Kramer adjustment was made to p-values to compensate for multiple follow-up comparisons. All p-values that are reported have been adjusted accordingly.

To report effect size, Cohens *f*^2^ for multilevel models was estimated from the mixed models calculated in SAS using the process described by Selya et al. (2012). In addition, Cohen’s *d*_*z*_ for within-subjects comparisons was calculated to report effect size for all follow-up comparisons calculated from the LSmeans procedure.

An exploratory analysis was performed to test for potential moderating effect of gender. To do this, the eight mixed models were repeated, but this time included gender as a fixed effect and the two-way interactions of gender and condition, and gender and trial-type were evaluated. These models also tested the three-way interaction between gender, trial-type and condition.

To aid with interpretability and visual comparisons, z-scores were calculated for each of the primary dependent variables of interest (accuracy, RT, N2 component amplitude, P3 component amplitude) for both the go/no-go task and the flanker task. N2 amplitude and RT values were reverse scored as a more negative N2 is seen as larger and faster response time is seen as improved performance. These relationships as a function of exercise condition are displayed in Figure 2.

**Figure 2:**
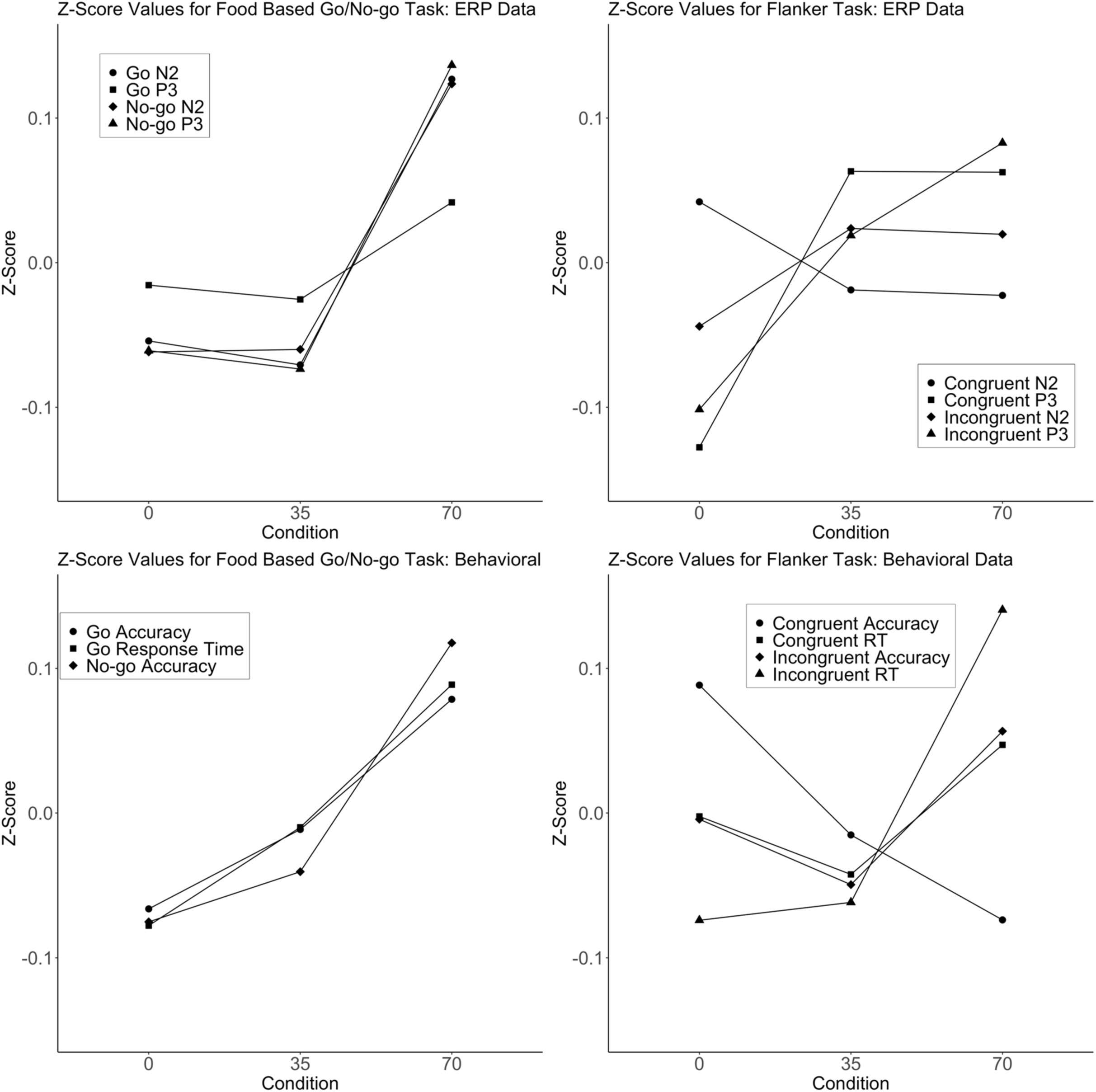
Z-score values of dependent variables for both tasks by exercise condition. Scores for negative-going measures (e.g., N2 amplitude, response times) were reversed so higher z-scores are associated with larger component amplitude or faster performance for ease of interpretation.

## 3. Results

### 3.1 Accuracy

Overall, accuracy on the go/no-go task was high for all conditions and trials (see Table 4). There was a significant main effect for condition (rest, 35% and 70%; *F*(2,420) = 4.56, *p* = .01, *f*^*2*^ = 0.01) and trial-type (go vs. no-go; *F*(1,636) = 776.08, *p* < .001, *f*^*2*^ = 0.59) but no significant interaction between condition and trial-type (*F*(2,636) = 0.30, *p* = .74, *f*^*2*^ < 0.01). Accuracy was better for the go trials compared to the no-go trials, as expected. Task accuracy was better for the 70% condition compared to the rest condition (*t*(2,420) = −3.01, *p* < .01, *d*_*z*_ = 0.15), but the 70% condition was not different compared to the 35% condition (*t*(2,420) = −1.35, *p* = .37, *d*_*z*_ = 0.09). There was also no difference between the 35% and rest conditions (*t*(2,420) = −1.66, *p* = .22, *d*_*z*_ = 0.07).

For the flanker task, participants were more accurate on the congruent trials compared to the incongruent trials (*F*(1,636) = 626.52, *p* < .001, *f*^*2*^ = 0.38; see Table 5), as expected. However, there was no main effect for exercise condition (*F*(2,420) = 1.83, *p* = .16, *f*^*2*^ < 0.01) along with no interaction for accuracy between condition and trial-type (*F*(2,636) = 1.13, *p* = .32, *f*^*2*^ < 0.01).

### 3.2 Response times

For the go/no-go task, correct go trial response time was different between the exercise conditions (*F*(2,420) = 6.52, *p* = .002, *f*^*2*^ = 0.03; see Table 4). Response times following the 70% condition were significantly faster than the rest condition (*t*(2,420) *=* 3.60, *p* < .001, *d*_*z*_ = 0.30) but were not different compared to the 35% condition (*t*(2,420) = 2.05, *p* = .10, *d*_*z*_ = 0.13). There was also no difference in response time between the rest and 35% conditions (*t*(2,420) = 1.54, *p* = .27, *d*_*z*_ = 0.12).

For the flanker task, there was a significant main effect of exercise condition (*F*(2,420) = 7.47, *p* < .001, *f*^*2*^ = 0.03) and trial-type (*F*(1,635) = 6216.94, *p* < .001, *f*^*2*^ = 3.27), along with a significant condition by trial-type interaction (*F*(2,635) = 5.28, *p* < .01, *f*^*2*^ = 0.02). Response times were faster for the congruent compared to incongruent trials, as expected. Response times were faster for the 70% condition compared to both the rest (*t*(2,420) = 2.95, *p* = .009, *d*_*z*_ *=* 0.14) and 35% conditions (*t*(2,420) = 3.64, *p* = .001, *d*_*z*_ *=* 0.17). There was no difference in response time between the rest and 35% conditions (*t*(2,420) = −0.70, *p* = 0.763, *d*_*z*_ *<* .01). Faster response times following the 70% condition compared to the other conditions were qualified by a significant condition by trial-type interaction, where the increase in response speed during the 70% condition was observed primarily during the incongruent compared to the congruent trials for both the rest (*t*(1,635) = 4.15, *p* < 0.001, *d*_*z*_ *=* 0.29) and 35% conditions (*t*(1,635) = 4.17, *p* < .001, *d*_*z*_ *=* 0.27; see Table 5).

### 3.3 Food related inhibitory control

Figure 3 displays the N2 and P3 waveforms by exercise condition for the food-based go/no-go task. There was a significant main effect for both trial-type (*F*(1,610) = 164.81, *p* < .001, *f*^*2*^ = 0.05; see Table 4) and exercise condition (*F*(2,394) = 4.63, *p* = .01, *f*^*2*^ = < 0.02) for the N2 component, however, there was no significant condition by trial-type interaction (*F*(2,610) = 0.09, *p* = .92, *f*^*2*^ < 0.01). The N2 amplitude was more negative (i.e., larger) for no-go trials than go trials, as expected. Follow-up analyses demonstrated that the N2 for the 70% condition was significantly more negative for both the go and the no-go trials compared to both the rest (*t*(2,394) = 2.46, *p* = 0.038, *d*_*z*_ = 0.17) and 35% conditions (*t*(2,394) = 2.79, *p* = 0.015, *d*_*z*_ = 0.18). There was no difference in N2 amplitude between the rest and 35% conditions (*t*(2, 394) = −0.31, *p* < 0.947, *d*_*z*_ = 0.02).

**Figure 3:**
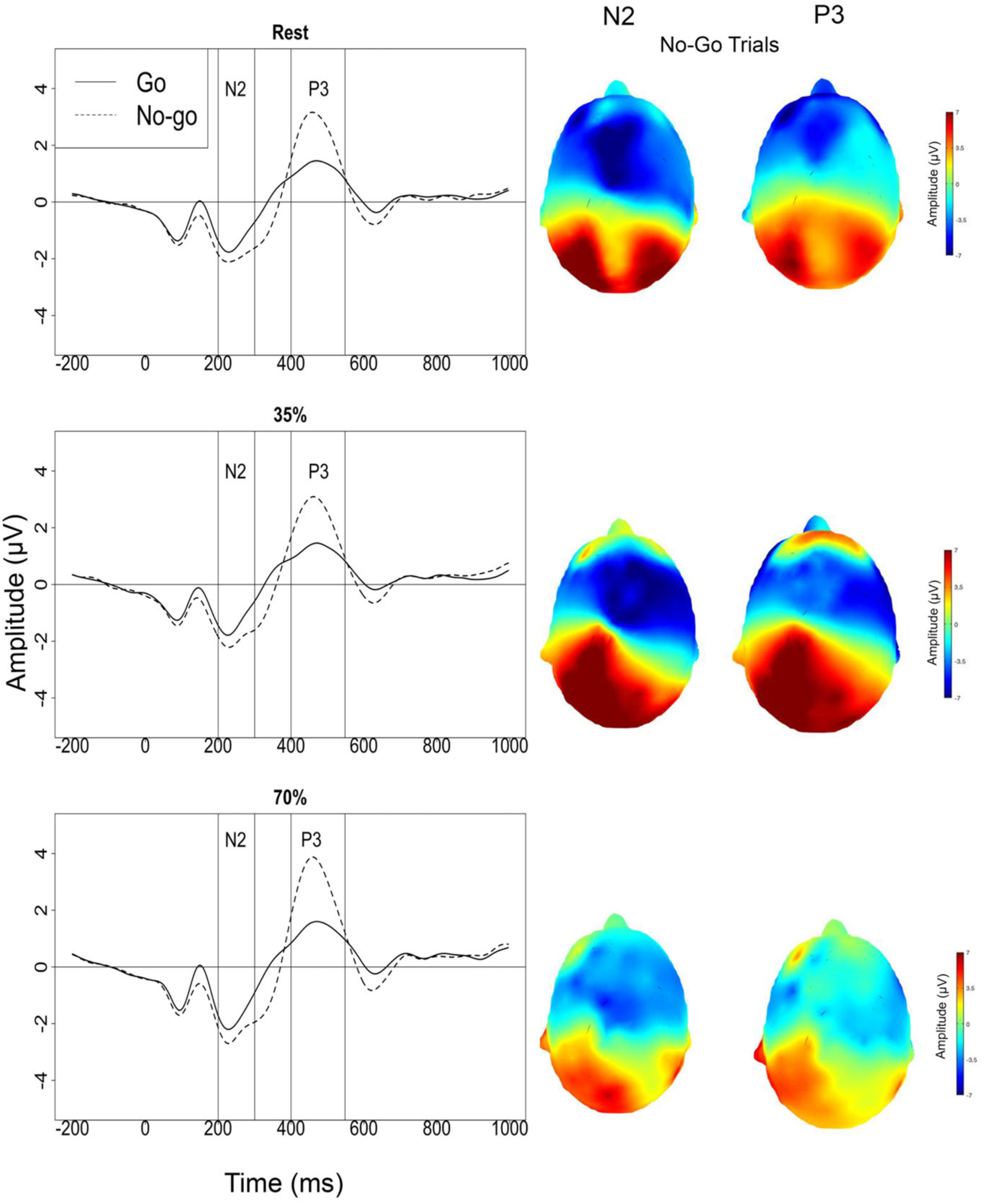
Event-related potential waveforms and topographical maps for the N2 and P3 on the go/no-go trials.

Similar to the N2, the P3 ERP component also demonstrated a significant main effect for both trial-type (*F*(1,610) = 432.45, *p* < 0.001, *f*^*2*^ = 0.29) and exercise condition (*F*(2,394) = 4.84, *p* = 0.008, *f*^*2*^ = 0.02; see Table 4). There was also a significant interaction between trial-type and condition (*F*(2,610) = 3.56, *p* = 0.03, *f*^*2*^ *=* 0.01). The no-go trials displayed a significantly more positive (i.e., larger) P3 amplitude than the go trials, as expected. The P3 for the go trials was not different between the three conditions (*p*’s > 0.05), however for the no-go trials the P3 was significantly more positive after the 70% condition compared to both the rest (*t*(1, 610) = −3.52, *p* = 0.006, *d*_*z*_ = 0.22) and 35% conditions (*t*(1, 610) = −3.53, *p* = 0.006, *d*_*z*_ = 0.24).

### 3.4 Cognitive control

Figures 4 and 5 display the N2 and P3 waveforms by exercise conditions for the flanker task. For the N2 component during the flanker task, there was a main effect of trial-type (congruent vs. incongruent; *F*(1,593) = 279.71, *p* < .001, *f*^*2*^ = 0.21; see Table 4) but no main effect of exercise condition (*F*(2,377) = 0.09, *p* = .91, *f*^*2*^ < 0.01) nor an interaction between condition and trial-type (*F*(2,593) = 0.82, *p* = .44, *f*^*2*^ < 0.01). The N2 for the incongruent trials was more negative when compared to the congruent trials, as expected.

**Figure 4:**
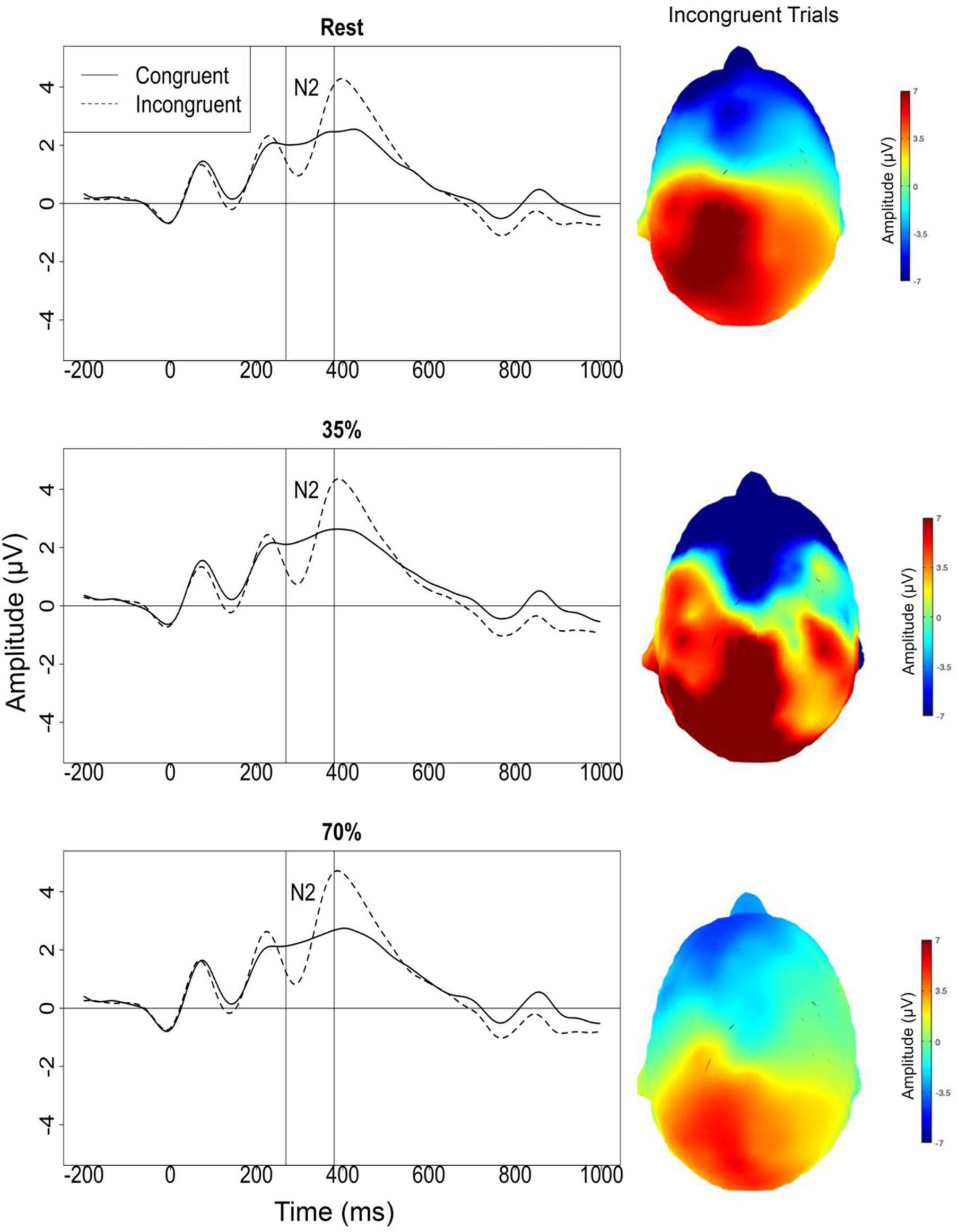
Event-related potential waveforms and topographical maps for the N2 on the incongruent trials during the flanker task.

**Figure 5:**
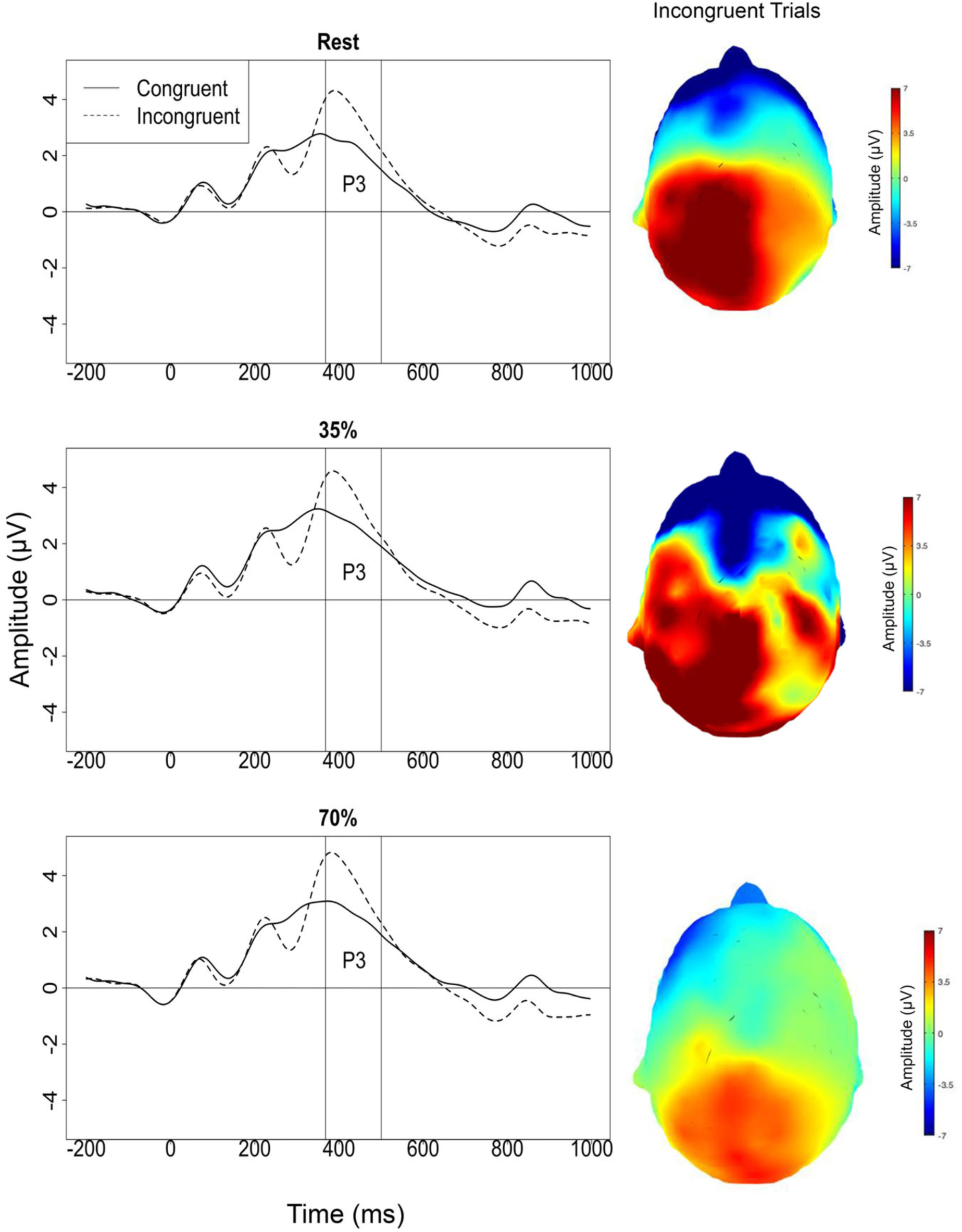
Event-related potential waveforms along with topographical maps for the P3 on the incongruent trials during the flanker task.

In contrast, for the P3 component there was a significant main effect for condition (*F*(2, 377) = 3.60, p = 0.03, *f*^*2*^ = 0.01) and trial-type (*F*(1,593) = 199.10, *p* < 0.001, *f*^*2*^ = 0.17) but there was no interaction between condition and trial-type (*F*(2,593) = 0.30, *p* = 0.739, *f*^*2*^ < 0.01). Incongruent trials elicited a more positive P3 response when compared to congruent trials, as expected. The 70% condition was significantly more positive than the rest condition (*t*(2, 377) = −2.60, *p* = .03, *d*_*z*_ = 0.17) but was not different than the 35% condition (*t*(2, 377) = −0.74, *p* = .74, *d*_*z*_ = 0.04). There was no significant difference between the rest and 35% conditions (*t*(2, 377) = − 1.90, *p* = .14, *d*_*z*_ = 0.14).

### 3.5 Gender

Including gender in the previous models had no impact on the interpretation of any of exercise condition-related relationships. In other words, there were no gender-by-condition interactions for any of the exercise condition analyses and there were no three-way interactions between gender, condition and trial-type for any of the primary dependent variables of interest (RT, accuracy, N2, and P3 components). There was a significant difference between genders for food go/no-go N2 (*F*(1, 606) = 5.39, *p* = .02), and flanker P3 (*F*(1, 589) = 6.66, *p* = .01), accuracy (*F*(1, 632) = 13.28, *p* < .001), and response time (*F*(1, 631) = 13.69, *p* < 0.001). Men had more negative go/no-go N2 amplitudes, more positive flanker P3 amplitudes, greater flanker accuracy and faster flanker response times.

There was also a significant gender-by-trial-type interaction for food go/no-go N2 (*F*(1, 606) = 6.43, *p* = .01), and flanker N2 (*F*(1, 589) = 20.11, *p* < .001), P3 (*F*(1, 589) = 6.32, *p* = .01), accuracy (*F*(1, 632) = 5.05, *p* = .02), and response time (*F*(1, 631) = 37.63, *p* < .001). The difference between go and no-go trial N2 amplitude was greater for men than women (0.22 ± 0.08 μV; *t*(2, 606) = 2.54, *p* = 0.011). The difference between the incongruent and congruent trials was greater for men than women for both the N2 (0.61 ± 0.14 μV; *t*(1, 589) = −4.48, p < 0.001) and P3 (0.49 ± 0.19 μV; *t*(1, 589) = 2.51, *p* = .012) ERP components. The difference in accuracy between the incongruent and congruent trials was greater in women than men (0.01 ± 0.006 %; *t*(1, 632) = 2.25, *p* = .025). Similarly, the difference in response time between the incongruent and congruent trials was greater for women than men (9.52 ± 1.55 ms; *t*(1, 631) = − 6.13, *p* < .001).

## 4. Discussion

We used a high-powered, within-subjects crossover design to test the role of exercise intensity on behavioral and neurophysiological measures of cognitive control (flanker task performance and ERP amplitudes) and food-related inhibitory control (go/no-go task performance and ERP amplitudes). The impact of exercise on cognitive control (as measured by the flanker task) was intensity dependent. Specifically, response times were faster following vigorous intensity exercise at 70% of max VO_2max_ compared to both rest and moderate intensity exercise at 35% of max VO_2max_ and P3 component amplitudes for congruent and incongruent trials were more positive following vigorous intensity exercise compared to rest, but not moderate intensity exercise. Notably, response times were disproportionately faster with higher intensity exercise on incongruent compared to congruent trials, suggesting that cognitive control may be specifically more efficient following high intensity exercise. N2 component amplitudes during the flanker task and flanker accuracy did not differ as a function of exercise intensity, indicating that the effects may not be present in all aspects of cognitive control performance, although nuance is required in the interpretation of these findings.

The impact of exercise on cognitive control has been evaluated in a number of studies, the majority of which have shown enhanced P3 component amplitude following exercise (Kao et al., 2019) with relatively few studies reporting N2 component results. For example, current N2 results that do not differ by exercise condition are consistent with Themanson et al. (2006) who also observed no change in N2 amplitude during a modified flanker task following exercise at a similar intensity as prescribed in our study (roughly 85% of heart rate max, or 169 beats per minute). Similarly, recent work by Chacko et al., (2020) suggests that vigorous-intensity aerobic exercise may be more related to selective attention, but not initial indexing of conflict or control- related functions—consistent with findings of enhanced P3 component amplitude, but no condition-related differences for N2 amplitude. However, current N2 results are in contrast to those observed by Ligeza et al. (2018) who observed a more negative N2 following exercise between the 1^st^ and 2^nd^ ventilatory thresholds, which turned out to be about 75% of heart rate max. They also observed a blunted (i.e., less negative) N2 following high intensity interval training.

Key differences in these studies may explain the seemingly divergent results. First, the exercise performed in our study was most similar to Themanson et al (2006) and in-between the intensities performed in the Ligeza et al. (2018) study. Specifically, our moderate exercise (35% VO_2max_) condition was less intense than the aerobic exercise prescribed in Ligeza et al. (2018), and our vigorous exercise condition (70%) was less intense than their high intensity interval training condition. In addition, each participant completed the flanker task after the food related go/no-go task, which may have influenced our results as the effects of exercise on brain activity may change with time (Ciria et al., 2018).

There are over 20 studies that have evaluated the impact of exercise on attentional allocation measured by P3, although it is challenging to bring the findings of these studies together given the variability in exercise duration and intensity, the timing of the neural measurement post-exercise, and the variety of different cognitive tasks performed (Kao et al., 2019; Ludyga et al., 2016). In addition, most of the studies had small sample sizes, which reduces statistical power and limits the ability to accurately identify small effects of exercise on cognitive control. The large sample and within-subjects design of the current research considerably increases the statistical power and confidence in current results, along with the consistent findings with the large majority of the cognitive control and P3 amplitude literature (Kao et al., 2019).

Specifically, studies evaluating exercise completed in the light- to low-moderate intensity range (similar to light walking) seem to agree with our finding that there is no impact on P3 amplitude (Kamijo et al., 2004; Kamijo and Takeda, 2009). However, studies that exercise participants at an intensity that is similar to 60 to 75% of heart rate max generally demonstrate that the P3 ERP component is elevated compared to controls (Chang et al., 2017; Kamijo et al., 2004; Kamijo et al., 2007; Kao et al., 2017; O’Leary et al., 2011; Pontifex et al., 2015; Scudder et al., 2012). The effect of exercise seems to weaken as intensity of exercise increases, with mixed results for studies exceeding 75% of heart rate max (Chu et al., 2015; Hillman et al., 2003; Kamijo et al., 2007; Ligeza et al., 2018). Very high intensity exercise of greater than 90% seems to either have no impact on the P3 ERP component or a decreased P3 amplitude suggesting that the inverted U we initially hypothesized may be present at higher intensities of exercise than was conducted in the current research (Kamijo et al., 2004; Kao et al., 2017; Ligeza et al., 2018). Thus, current results are a step in understanding the role of exercise intensity levels on cognitive control functions, but future studies testing higher intensity levels are needed to more fully test an inverted-U hypothesis.

A critical component of cognitive control is response inhibition, which involves overcoming or suppression of an action that is inappropriate in a given context. For the current study we were interested in the response to high- and low-calorie foods tested using a food-specific go/no-go task. A clear pattern of improved performance and increased ERP amplitudes was present for vigorous exercise (70% of VO_2max_) compared to rest. Specifically, there was improved accuracy, faster response times, and larger N2 and P3 amplitudes for vigorous exercise. Notably, these were all main effects with the exception of P3 component amplitude that showed an interaction and was specific to no-go, but not go trials. These results suggest that the impact of exercise on food-specific response inhibition may be a more general facilitative affect and that this facilitative effect is intensity dependent. Both the N2 and P3 were larger in the vigorous exercise condition compared to the moderate (35% of VO_2max_) exercise condition and the seated rest condition. Response time and accuracy were both better for the vigorous exercise condition compared to the rest condition.

Notably, there were no differences between the moderate intensity exercise and rest conditions. The absence of condition-related differences between the moderate intensity exercise and rest conditions is a consistent finding for both the food-specific go/no-go and flanker tasks— indicating that higher intensity exercise appears necessary to modulate inhibitory and cognitive control measures. More specifically, although 35% of VO_2max_ is classified as moderately intense exercise (3-6 METs), these results suggest that 35% of VO_2max_ is insufficiently intense to have a meaningful impact on subsequent neural activity during the go/no-go task. The 70% condition is a vigorously intense activity level (> 6 METS) and while we anticipated a suppression of neural activity at this intensity, the results of the study suggest that if there is a U-shaped relationship, 70% of VO_2max_ is still in the range where neural activity is elevated.

In general, the findings from the go/no-go food-specific inhibitory control task largely parallel the changes in general cognitive control as observed from the flanker task. Two main exceptions to this that may point to a food specific effect of exercise on response inhibition. First, there was a significant main effect of condition on the N2 component during the go/no-go task, but no main effect of condition during the flanker task. Second, there was a trial-type-by-condition interaction for the P3 ERP component during the go/no-go task, but this same interaction was not observed during the flanker task. During the go/no-go task a more positive P3 result was observed specifically in the 70% condition for no-go high-calorie pictures, suggesting increased recruitment of later neural resources to increase inhibitory control specifically towards high-calorie foods. This elevated P3 component during no-go trials following vigorous exercise suggests that more neural resources were recruited to inhibit the dominant response for high-calorie foods.

There are only a handful of studies that have evaluated the impact of exercise on various event-related potentials to visual food cues. These studies have primarily used passive viewing tasks in contrast to cognitive control tasks (such as a go/no-go task). For example, Hanlon et al., (2012) showed a reduced late posterior positivity (LPP) amplitude to pictures of plated foods compared to pictures of flowers in women (both healthy weight and those with obesity) after an acute 45-minute bout of high moderate-intensity exercise compared to rest, suggesting reduced motivation towards food following exercise. In contrast, participants in Carbine et al. (In Press) performed exercise at moderate (3.7 METS) and vigorous (7.4 METS) intensities and found no difference in the centro-parietal P3 or LPP ERP components following either exercise condition or rest. Given that these two studies were conducted using passive viewing attention-based paradigms, instead of cognitive control or response inhibition paradigms, it is difficult to make clear comparisons.

There are some studies in adolescents looking at exercise and neural responses to food cues. Two separate studies of adolescents showed decreased P3 amplitudes to food stimuli compared to nonfood stimuli following acute moderate-intensity exercise compared to rest (Fearnbach et al., 2016; Fearnbach et al., 2017). However, the decreased amplitudes were moderated by obesity status, since the P3 component amplitude to food stimuli was decreased after exercise only in adolescents with obesity. In another crossover study in adolescents there was no difference in N2 amplitude to a go/no-go task between 60 minutes of seated video game play vs. active video game play (Smith et al., 2020). Taken together, all these studies suggest that there might be a positive role of exercise on reducing attentional allocation towards food cues, however the results are not homogeneous and may be more impactful in adults or be moderated by obesity. Our study adds to this research, suggesting that not only does exercise have the potential to influence neural reflections on food cues but also is able to enhance response inhibition to high-calorie foods. Additionally, our study adds evidence that primarily vigorous exercise (70% VO_2max_), rather than moderate exercise (35% VO_2max_), may be beneficial in increasing cognitive control and food-specific inhibitory control functions.

As an exploratory portion of this study, we tested the possible moderating role of gender. The gender-related analyses were done primarily to inform future research and to ensure that there were not gender-specific effects given recent findings suggesting that P3 component amplitude to a flanker task may be more positive in female than male exercisers, but not different between genders in more sedentary individuals (Lennox et al., 2019). Notably, we did not have gender-related hypotheses going into the current study. In contrast to the Lennox et al. findings, gender did not interact with exercise intensity for any of the dependent variables in the current study. Despite the absence of interactions with exercise intensity, there were some gender differences that are worth noting even though they did not alter the results of this study. Men in the study tended to have a more negative N2 response for the go/no-go task and a more positive P3 response to the flanker task. They also tended to respond faster and with better accuracy than the women. Our findings of increased amplitude N2 and P3 component amplitudes in men compared to women are consistent with previous work showing larger N2 and error-related ERP component amplitudes in men compared to women as well as decreased accuracy and longer response times in female participants (Clayson et al., 2011; Larson et al., 2011b; Stoet, 2010), suggesting these gender and cognitive control differences are likely not specific to exercise.

### 4.1 Potential mechanisms

Although there have been a number of proposed mechanisms explaining the impact of exercise on response inhibition and cognitive control, the mechanisms underlying these changes remain unclear and evidence is still limited. One possible mechanism is neurotransmitter changes associated with exercise—specifically catecholamines that are associated with cognitive response following higher intensity exercise (e.g., exercise beyond moderate walking)(Joris et al., 2018; Ogoh and Ainslie, 2009). The possibility of catecholamine-related changes is also interesting since catecholamines are involved in altering eating behavior (Wellman, 2000). The concentration of catecholamines in the brain raises with increased exercise intensity but appreciable levels of norepinephrine are not generally observed until around 50% of VO_2max_ (Joris et al., 2018; Ogoh and Ainslie, 2009). This would potentially explain the increase in ERP amplitudes, accuracy and response times in the vigorous intensity condition compared to both the moderate and rest conditions. It is also possible that elevated levels of serum brain derived neurotrophic factor (BDNF) can modulate the relationship between exercise intensity and cognitive performance, particularly since BDNF increases at moderate to high levels of physical activity (Hung et al., 2018; Jimenez-Maldonado et al., 2018).

Another possible moderator of the relationship between exercise intensity and cognitive and inhibitory control functions is changes in cerebral blood flow following exercise (Smith and Ainslie, 2017). Specifically, there is a transient change in cerebral blood flow that is intensity dependent. As exercise becomes more intense, cerebral blood flow tends to increase up to an exercise intensity of ∼60% VO_2max_ after which blood flow plateaus and begins to decrease toward resting values as exercise intensity continues to increase, likely due to vasoconstriction (Joris et al., 2018; Ogoh and Ainslie, 2009; Smith and Ainslie, 2017). As noted above, it may be that the current ∼70% VO_2max_ is not sufficiently vigorous to see a downturn in performance. Future research is needed to address this possibility.

There are other neuroelectric mechanisms specific to ERPs and human electrophysiology that may explain some variance in the relationship between exercise intensity and cognitive/inhibitory control abilities. Specifically, Polich (2007) suggested that increased P3 amplitude was the neural response stemming from memory processing coming from inhibiting task-irrelevant brain activation. This inhibition may be influenced by exercise-related changes in arousal regulated by the reticular activating system (Kinomura et al., 1996; McMorris et al., 2018), although recent findings suggest that the locus coeruleus may not be specifically implicated (McGowan et al., 2019). Others suggest that an increase in P3 amplitude may be due to a general arousal effect associated with exercise-related activity that heightens neuroelectric activity (Magnie et al., 2000). In short, although the precise mechanisms remain nonspecific, current findings likely result from an interaction of neurotransmitter, hemodynamic, and neuroelectric increases that may be associated with the general arousal following high intensity exercise.

### 4.2 Study limitations and strengths

Study limitations should be considered when interpreting the current results. First, the testing order for the food go/no-go and the flanker was not randomized. The study was specifically designed for the food go/no-go paradigm to be completed first since food-related inhibitory control was the primary research question and the general cognitive control question was secondary. Not controlling for task order in the design means we cannot rule-out the possibility that the flanker results were influenced by the food-specific go/no-go task. Differences in findings between the tasks may be related to the timing of presentations, since the flanker was always performed after the go/no-go task. In addition, while the gender differences were interesting, the sampling for the study was not done randomly. Thus, the interpretation of differences between genders should be considered with caution.

Despite the limitations, this study is unique and has several strengths. First, this study is one of the first to evaluate the impact of acute exercise across multiple intensities on food-related response inhibition. In addition, the large sample size makes the estimates of effect size more stable and reduces the possibility of inflated effects or missing a small effect. In addition, the study included both genders in roughly equal number, which allowed for some inferences on the role of gender on these relationships. Previous investigations lacked the sample size and gender diversity to explore this question. While not perfectly designed (as go/no-go and flanker task order was not randomized) the study also attempted to evaluate if any changes in food-related neural inhibition were a result of general cognitive control changes. The study also is unique in that it evaluates two different exercise intensities that are commonly performed at a duration that is more consistent with weight management recommendations. The apparent differences in the neural and behavioral response between exercise intensities suggests that evaluating different exercise intensities is important because the results of the study change based on the exercise prescription. Finally, the exercise prescription was precisely prescribed base on individually-measured maximal aerobic capacity, which is different for each individual.

## 5 Conclusions and future directions

Overall, this study supports previous studies that a single bout of exercise has the potential to influence measures of food-related inhibitory control and cognitive control. Results suggest that these benefits extend primarily to higher (i.e., jogging) but not lower intensity exercise (i.e., light walking). Benefits to inhibitory control are important for daily tasks such as withholding a prepotent response (like eating an apple instead of a donut when both are available) particularly during nonroutine circumstances (Banich, 2009). Thus, higher intensity aerobic exercise seems to be an efficient means of increasing inhibitory and cognitive control functions for a period of time after exercise (Kao et al., 2019). Future research is needed to determine specific neural mechanisms, assess if there is an inverted-U at maximal intensity thresholds, test the role of adiposity/obesity, and to evaluate how food-specific inhibitory control and cognitive control are altered with exercise training over time and individual aerobic fitness.

## Acknowledgements

The current study required a large team effort. We acknowledge and thank the following for their contributions to the project: Whitney D. Allen, Erin M. Corbin, Emma Hedges, Hanel Watkins, Jase Hoskin, Mika Honda, Emma Gleave, Michael Christensen, Landon Deru, Rebekah E. Rodeback, Tanner Ward, Landon Johnson, Joseph Hicks, Ryland Savage, Matt Starr, Andrew Stevens, Caleb Summerhays, Timothy Swingle, Kekoa Wu, James Whitlock, Hiilei Ellis, and Mitch Rands.

## Author Contributions

Bruce Bailey: conceptualization, methodology, formal analysis, resources, data curation, writing – original draft, visualization, supervision, project administration, funding acquisition. Alexandra M. Muir: data acquisition, visualization, formal data analysis, writing the original manuscript draft, writing – original draft, writing - review & editing, data curation. Ciera Bartholomew: data acquisition, visualization, formal data analysis, writing – original draft, writing - review & editing. William F. Christensen: formal data analysis, writing - review & editing. Kaylie A. Carbine: supervision, data curation, writing - review & editing. Harrison Marsh: supervision, project administration, writing - review & editing. Hunter LaCouture: supervision, project 733 administration, writing - review & editing. Chance McCutcheon: data curation, writing – review 734 & editing. Michael J. Larson: conceptualization, funding acquisition, formal data analysis, 735 methodology, resources, visualization, writing – original draft, writing - review & editing.

## References

ACSM, 2018. ACSM’s Guidelines for Exercise Testing and Prescription: 10thEdition. Baltimore, Maryland. Lippincott Williams & Wilkins.

Allom, V., Mullan, B., 2014. Individual differences in executive function predict distinct eating behaviours. Appetite 80, 123–130. https://doi.org/10.1016/j.appet.2014.05.007

Armelagos, G.J., 2014. Brain evolution, the determinates of food choice, and the omnivore’s dilemma. Crit Rev Food Sci Nutr 54, 1330–1341. https://doi.org/10.1080/10408398.2011.635817

Arraiz, G.A., Wigle, D.T., Mao, Y., 1992. Risk assessment of physical activity and physical fitness in the Canada Health Survey mortality follow-up study. J Clin Epidemiol 45, 419–428. https://doi.org/10.1016/0895-4356(92)90043-m

Aulbach, M.B., Harjunen, V.J., Spapé, M., Knittle, K., Haukkala, A., Ravaja, N., 2020. No evidence of calorie-related modulation of N2 in food-related Go/No-Go training: A preregistered ERP study. Psychophysiology 57, e13518. https://doi.org/10.1111/psyp.13518

Banich, M.T., 2009. Executive function: the search for an integrated account. Curr. Dir. Psychol. Sci. 18, 89–94. https://doi.org/10.1111/j.1467-8721.2009.01615.x

Botvinick, M.M., Braver, T.S., Barch, D.M., Carter, C.S., Cohen, J.D., 2001. Conflict monitoring and cognitive control. Psychol. Rev. 108, 624–652. https://doi.org/10.1037/0033-295x.108.3.624

Brush, C.J., Bocchine, A.J., Olson, R.L., Ude, A.A., Dhillon, S.K., Alderman, B.L., 2020. Does aerobic fitness moderate age-related cognitive slowing? Evidence from the P3 and lateralized readiness potentials. Int. J. Psychophysiol. 155, 63–71. http://doi.org/10.1016/j.ijpsycho.2020.05.007

Byun, K., Hyodo, K., Suwabe, K., Ochi, G., Sakairi, Y., Kato, M., Dan, I., Soya, H., 2014. Positive effect of acute mild exercise on executive function via arousal-related prefrontal activations: an fNIRS study. NeuroImage 98, 336–345. https://doi.org/10.1016/j.neuroimage.2014.04.067

Carbine, K.A., Anderson, J., Larson, M. J., LeCheminant, J. D., Bailey, B., In Press. The relationship between exercise intensity and neurophysiological responses to food stimuli in women: a randomized crossover event-related potential (ERP) study. Int. J. Psychophysiol. https://doi.org/10.31219/osf.io/k26md

Carbine, K.A., Christensen, E., LeCheminant, J.D., Bailey, B.W., Tucker, L.A., Larson, M.J., 2017. Testing food-related inhibitory control to high- and low-calorie food stimuli: Electrophysiological responses to high-calorie food stimuli predict calorie and carbohydrate intake. Psychophysiology 54, 982–997. https://doi.org/10.1111/psyp.12860

Carbine, K.A., Duraccio, K.M., Kirwan, C.B., Muncy, N.M., LeCheminant, J.D., Larson, M.J., 2018a. A direct comparison between ERP and fMRI measurements of food-related inhibitory control: Implications for BMI status and dietary intake. NeuroImage 166, 335–348. https://doi.org/10.1016/j.neuroimage.2017.11.008

Carbine, K.A., Rodeback, R., Modersitzki, E., Miner, M., LeCheminant, J.D., Larson, M.J., 2018b. The utility of event-related potentials (ERPs) in understanding food-related cognition: A systematic review and recommendations. Appetite 128, 58–78. https://doi.org/10.1016/j.appet.2018.05.135

Carek, P.J., Laibstain, S.E., Carek, S.M., 2011. Exercise for the treatment of depression and anxiety. Int. J. Psychiatry Med. 41, 15–28. https://doi.org/10.2190/PM.41.1.c

Chacko, S.C., Quinzi, F., De Fano, A., Bianco, V., Mussini, E., Berchicci, M., Perri, R.L., Di Russo, F., 2020. A single bout of vigorous-intensity aerobic exercise affects reactive, but not proactive cognitive brain functions. Int. J. Psychophysiol. 147, 233–243. http://doi.org/10.1016/j.ijpsycho.2019.12.003

Chang, Y.K., Chu, I.H., Liu, J.H., Wu, C.H., Chu, C.H., Yang, K.T., Chen, A.G., 2017. Exercise modality is differentially associated with neurocognition in older adults. Neural Plast. https://doi.org/10.1155/2017/3480413

Chang, Y.K., Labban, J.D., Gapin, J.I., Etnier, J.L., 2012. The effects of acute exercise on cognitive performance: a meta-analysis. Brain research 1453, 87–101. https://doi.org/10.1016/j.brainres.2012.02.068

Chu, C.-H., Alderman, B.L., Wei, G.-X., Chang, Y.-K., 2015. Effects of acute aerobic exercise on motor response inhibition: An ERP study using the stop-signal task. J Sport Health Sci 4, 73–81. https://doi.org/10.1016/j.jshs.2014.12.002

Ciria, L.F., Perakakis, P., Luque-Casado, A., Sanabria, D., 2018. Physical exercise increases overall brain oscillatory activity but does not influence inhibitory control in young adults. NeuroImage 181, 203–210. https://doi.org/10.1016/j.neuroimage.2018.07.009

Clayson, P.E., 2020. Moderators of the internal consistency of error-related negativity scores: A meta-analysis of internal consistency estimates. Psychophysiology 57, e13583. http://doi.org/10.1111/psyp.13583

Clayson, P.E., Baldwin, S.A., Larson, M.J., 2013. How does noise affect amplitude and latency measurement of event-related potentials (ERPs)? A methodological critique and simulation study. Psychophysiology 50, 174–186. https://doi.org/10.1111/psyp.12001

Clayson, P.E., Carbine, K.A., Baldwin, S.A., Larson, M.J., 2019. Methodological reporting behavior, sample sizes, and statistical power in studies of event-related potentials: Barriers to reproducibility and replicability. Psychophysiology 56, e13437. http://doi.org/10.1111/psyp.13437

Clayson, P.E., Clawson, A., Larson, M.J., 2011. Sex differences in electrophysiological indices of conflict monitoring. Biol Psychol 87, 282–289. http://doi.org/10.1016/j.biopsycho.2011.03.011

Clayson, P.E., Miller, G.A., 2017. ERP Reliability Analysis (ERA) Toolbox: An open-source toolbox for analyzing the reliability of event-related brain potentials. Int. J. Psychophysiol. 111, 68–79. https://doi.org/10.1016/j.ijpsycho.2016.10.012

Cornelissen, V.A., Smart, N.A., 2013. Exercise training for blood pressure: a systematic review and meta-analysis. J Am Heart Assoc 2, e004473. https://doi.org/10.1161/jaha.112.004473

Diamond, A., 2013. Executive functions. Annu Rev Psychol 64, 135–168. https://doi.org/10.1146/annurev-psych-113011-143750

Dien, J., 2010. The ERP PCA Toolkit: an open source program for advanced statistical analysis of event-related potential data. Journal of neuroscience methods 187, 138–145. https://doi.org/10.1016/j.jneumeth.2009.12.009

Drollette, E.S., Scudder, M.R., Raine, L.B., Moore, R.D., Saliba, B.J., Pontifex, M.B., Hillman, C.H., 2014. Acute exercise facilitates brain function and cognition in children who need it most: An ERP study of individual differences in inhibitory control capacity. Dev Cogn Neuros-Neth 7, 53–64. https://doi.org/10.1016/j.dcn.2013.11.001

Eriksen, B.A., Eriksen, C.W., 1974. Effects of noise letters upon the identification of a target letter in a nonsearch task. Perception & Psychophysics 16, 143–149. http://wexler.free.fr/library/files/eriksen%20(1974)%20effects%20of%20noise%20letters%20upon%20the%20identification%20of%20a%20target%20letter%20in%20a%20nonsearch%20task.pdf

Falkenstein, M., Hoormann, J., Hohnsbein, J., 1999. ERP components in Go/Nogo tasks and their relation to inhibition. Acta Psychol (Amst) 101, 267–291. https://doi.org/10.1016/s0001-6918(99)00008-6

Fearnbach, S.N., Silvert, L., Keller, K.L., Genin, P.M., Morio, B., Pereira, B., Duclos, M., Boirie, Y., Thivel, D., 2016. Reduced neural response to food cues following exercise is accompanied by decreased energy intake in obese adolescents. Int. J. Obes. 40, 77–83. https://doi.org/10.1038/ijo.2015.215

Fearnbach, S.N., Silvert, L., Pereira, B., Boirie, Y., Duclos, M., Keller, K.L., Thivel, D., 2017. Reduced neural responses to food cues might contribute to the anorexigenic effect of acute exercise observed in obese but not lean adolescents. Nutr Res 44, 76–84. https://doi.org/10.1016/j.nutres.2017.06.006

Fiuza-Luces, C., Santos-Lozano, A., Joyner, M., Carrera-Bastos, P., Picazo, O., Zugaza, J.L., Izquierdo, M., Ruilope, L.M., Lucia, A., 2018. Exercise benefits in cardiovascular disease: beyond attenuation of traditional risk factors. Nat. Rev. Cardiol. 15, 731–743. https://doi.org/10.1038/s41569-018-0065-1

Folstein, J.R., Van Petten, C., 2008. Influence of cognitive control and mismatch on the N2 component of the ERP: a review. Psychophysiology 45, 152–170. https://doi.org/10.1111/j.1469-8986.2007.00602.x

Gajewski, P.D., Falkenstein, M., 2013. Effects of task complexity on ERP components in Go/Nogo tasks. Int. J. Psychophysiol. 87, 273–278. https://doi.org/10.1016/j.ijpsycho.2012.08.007

George, J.D., 1996. Alternative approach to maximal exercise testing and VO2 max prediction in college students. Res Q Exerc Sport 67, 452–457. https://doi.org/10.1080/02701367.1996.10607977

Guerrieri, R., Nederkoorn, C., Jansen, A., 2007. How impulsiveness and variety influence food intake in a sample of healthy women. Appetite 48, 119–122. https://doi.org/10.1016/j.appet.2006.06.004

Guiney, H., Machado, L., 2013. Benefits of regular aerobic exercise for executive functioning in healthy populations. Psychon B Rev 20, 73–86. https://doi.org/10.3758/s13423-012-0345-4

Hagobian, T.A., Yamashiro, M., Hinkel-Lipsker, J., Streder, K., Evero, N., Hackney, T., 2013. Effects of acute exercise on appetite hormones and ad libitum energy intake in men and women. Appl. Physiol. Nutr. Metab. 38, 66–72. https://doi.org/10.1139/apnm-2012-0104

Hanlon, B., Larson, M.J., Bailey, B.W., LeCheminant, J.D., 2012. Neural response to pictures of food after exercise in normal-weight and obese women. Med Sci Sports Exerc 44, 1864–1870. https://doi.org/10.1249/MSS.0b013e31825cade5

Hillman, C.H., Snook, E.M., Jerome, G.J., 2003. Acute cardiovascular exercise and executive control function. Int. J. Psychophysiol. 48, 307–314. https://doi.org/10.1016/S0167-8760(03)00080-1

Hung, C.L., Tseng, J.W., Chao, H.H., Hung, T.M., Wang, H.S., 2018. Effect of Acute Exercise Mode on Serum Brain-Derived Neurotrophic Factor (BDNF) and Task Switching Performance. J Clin Med 7. https://doi.org/10.3390/jcm7100301

Jimenez-Maldonado, A., Renteria, I., Garcia-Suarez, P.C., Moncada-Jimenez, J., Freire-Royes, L.F., 2018. The Impact of High-Intensity Interval Training on Brain Derived Neurotrophic Factor in Brain: A Mini-Review. Front. Neurosci. 12. https://doi.org/10.3389/fnins.2018.00839

Joris, P.J., Mensink, R.P., Adam, T.C., Liu, T.T., 2018. Cerebral Blood Flow Measurements in Adults: A Review on the Effects of Dietary Factors and Exercise. Nutrients 10. https://doi.org/10.3390/nu10050530

Joseph, R.J., Alonso-Alonso, M., Bond, D.S., Pascual-Leone, A., Blackburn, G.L., 2011. The neurocognitive connection between physical activity and eating behaviour. Obes Rev 12, 800–812. https://doi.org/10.1111/j.1467-789X.2011.00893.x

Kamijo, K., Nishihira, Y., Hatta, A., Kaneda, T., Kida, T., Higashiura, T., Kuroiwa, K., 2004. Changes in arousal level by differential exercise intensity. Clin Neurophysiol 115, 2693–2698. https://doi.org/10.1016/j.clinph.2004.06.016

Kamijo, K., Nishihira, Y., Higashiura, T., Kuroiwa, K., 2007. The interactive effect of exercise intensity and task difficulty on human cognitive processing. Int. J. Psychophysiol. 65, 114–121. https://doi.org/10.1016/j.ijpsycho.2007.04.001

Kamijo, K., Takeda, Y., 2009. General physical activity levels influence positive and negative priming effects in young adults. Clin Neurophysiol 120, 511–519. https://doi.org/10.1016/j.clinph.2008.11.022

Kao, S.C., Cadenas-Sanchez, C., Shigeta, T.T., Walk, A.M., Chang, Y.K., Pontifex, M.B., Hillman, C.H., 2019. A systematic review of physical activity and cardiorespiratory fitness on P3b. Psychophysiology, e13425. https://doi.org/10.1111/psyp.13425

Kao, S.C., Cadenas-Sanchez, C., Shigeta, T.T., Walk, A.M., Chang, Y.K., Pontifex, M.B., Hillman, C.H., 2020. A systematic review of physical activity and cardiorespiratory fitness on P3b. Psychophysiology 57, e13425. https://doi.org/10.1111/psyp.13425

Kao, S.C., Westfall, D.R., Soneson, J., Gurd, B., Hillman, C.H., 2017. Comparison of the acute effects of high-intensity interval training and continuous aerobic walking on inhibitory control. Psychophysiology 54, 1335–1345. https://doi.org/10.1111/psyp.12889

Keil, A., Debener, S., Gratton, G., Junghofer, M., Kappenman, E.S., Luck, S.J., Luu, P., Miller, G.A., Yee, C.M., 2014. Committee report: Publication guidelines and recommendations for studies using electroencephalography and magnetoencephalography. Psychophysiology 51, 1–21. http://doi.org/10.1111/psyp.12147

Kempton, M.J., Ettinger, U., Foster, R., Williams, S.C., Calvert, G.A., Hampshire, A., Zelaya, F.O., O’Gorman, R.L., McMorris, T., Owen, A.M., Smith, M.S., 2011. Dehydration affects brain structure and function in healthy adolescents. Hum. Brain Mapp. 32, 71–79. https://doi.org/10.1002/hbm.20999

Killgore, W.D., Kipman, M., Schwab, Z.J., Tkachenko, O., Preer, L., Gogel, H., Bark, J.S., Mundy, E.A., Olson, E.A., Weber, M., 2013. Physical exercise and brain responses to images of high-calorie food. Neuroreport 24, 962–967. https://doi.org/10.1097/WNR.0000000000000029

Killgore, W.D., Young, A.D., Femia, L.A., Bogorodzki, P., Rogowska, J., Yurgelun-Todd, D.A., 2003. Cortical and limbic activation during viewing of high-versus low-calorie foods. NeuroImage 19, 1381–1394. https://doi.org/10.1016/s1053-8119(03)00191-5

Killgore, W.D., Yurgelun-Todd, D.A., 2005. Body mass predicts orbitofrontal activity during visual presentations of high-calorie foods. Neuroreport 16, 859–863. https://doi.org/10.1097/00001756-200505310-00016

Killgore, W.D., Yurgelun-Todd, D.A., 2007. Positive affect modulates activity in the visual cortex to images of high calorie foods. Int J Neurosci 117, 643–653. https://doi.org/10.1080/00207450600773848

Kinomura, S., Larsson, J., Gulyas, B., Roland, P.E., 1996. Activation by attention of the human reticular formation and thalamic intralaminar nuclei. Science 271, 512–515. https://doi.org/10.1126/science.271.5248.512

Kirwan, J.P., Sacks, J., Nieuwoudt, S., 2017. The essential role of exercise in the management of type 2 diabetes. Cleve. Clin. J. Med. 84, S15–s21. https://doi.org/10.3949/ccjm.84.s1.03

Knapen, J., Vancampfort, D., Moriën, Y., Marchal, Y., 2015. Exercise therapy improves both mental and physical health in patients with major depression. Disabil. Rehabil. 37, 1490–1495. https://doi.org/10.3109/09638288.2014.972579

Ko, Y.T., Miller, J., 2013. Signal-related contributions to stopping-interference effects in selective response inhibition. Exp Brain Res 228, 205–212. https://doi.org/10.1007/s00221-013-3552-y

Lambourne, K., Tomporowski, P., 2010. The effect of exercise-induced arousal on cognitive task performance: a meta-regression analysis. Brain research 1341, 12–24. https://doi.org/10.1016/j.brainres.2010.03.091

Larson, M.J., Carbine, K.A., 2017. Sample size calculations in human electrophysiology (EEG and ERP) studies: A systematic review and recommendations for increased rigor. Int. J. Psychophysiol. 111, 33–41. http://doi.org/10.1016/j.ijpsycho.2016.06.015

Larson, M.J., Clayson, P.E., Clawson, A., 2014. Making sense of all the conflict: a theoretical review and critique of conflict-related ERPs. Int. J. Psychophysiol. 93, 283–297. http://doi.org/10.1016/j.ijpsycho.2014.06.007

Larson, M.J., Farrer, T.J., Clayson, P.E., 2011a. Cognitive control in mild traumatic brain injury: conflict monitoring and conflict adaptation. Int. J. Psychophysiol. 82, 69–78. https://doi.org/10.1016/j.ijpsycho.2011.02.018

Larson, M.J., South, M., Clayson, P.E., 2011b. Sex differences in error-related performance monitoring. Neuroreport 22, 44–48. http://doi.org/10.1097/WNR.0b013e3283427403

Lavagnino, L., Arnone, D., Cao, B., Soares, J.C., Selvaraj, S., 2016. Inhibitory control in obesity and binge eating disorder: A systematic review and meta-analysis of neurocognitive and neuroimaging studies. Neurosci Biobehav Rev 68, 714–726. https://doi.org/10.1016/j.neubiorev.2016.06.041

LeBouthillier, D.M., Asmundson, G.J.G., 2017. The efficacy of aerobic exercise and resistance training as transdiagnostic interventions for anxiety-related disorders and constructs: A randomized controlled trial. J. Anxiety Disord. 52, 43–52. https://doi.org/10.1016/j.janxdis.2017.09.005

Lennox, K., Miller, R.K., Martin, F.H., 2019. Habitual exercise affects inhibitory processing in young and middle age men and women. Int. J. Psychophysiol. 146, 73–84. http://doi.org/10.1016/j.ijpsycho.2019.08.014

Ligeza, T.S., Maciejczyk, M., Kalamala, P., Szygula, Z., Wyczesany, M., 2018. Moderate-intensity exercise boosts the N2 neural inhibition marker: A randomized and counterbalanced ERP study with precisely controlled exercise intensity. Biol Psychol 135, 170–179. https://doi.org/10.1016/j.biopsycho.2018.04.003

Lowe, C.J., Kolev, D., Hall, P.A., 2016. An exploration of exercise-induced cognitive enhancement and transfer effects to dietary self-control. Brain Cogn 110, 102–111. https://doi.org/10.1016/j.bandc.2016.04.008

Luck, S.J., Gaspelin, N., 2017. How to get statistically significant effects in any ERP experiment (and why you shouldn’t). Psychophysiology 54, 146–157. https://doi.org/10.1111/psyp.12639

Ludyga, S., Gerber, M., Brand, S., Holsboer-Trachsler, E., Puhse, U., 2016. Acute effects of moderate aerobic exercise on specific aspects of executive function in different age and fitness groups: A meta-analysis. Psychophysiology 53, 1611–1626. https://doi.org/10.1111/psyp.12736

Mackie, M.A., Van Dam, N.T., Fan, J., 2013. Cognitive control and attentional functions. Brain Cogn 82, 301–312. https://doi.org/10.1016/j.bandc.2013.05.004

Magnie, M.N., Bermon, S., Martin, F., Madany-Lounis, M., Suisse, G., Muhammad, W., Dolisi, C., 2000. P300, N400, aerobic fitness, and maximal aerobic exercise. Psychophysiology 37, 369–377. https://doi.org/10.1017/S0048577200981435

McGowan, A.L., Chandler, M.C., Brascamp, J.W., Pontifex, M.B., 2019. Pupillometric indices of locus-coeruleus activation are not modulated following single bouts of exercise. Int. J. Psychophysiol. 140, 41–52. https://doi.org/10.1016/j.ijpsycho.2019.04.004

McMorris, T., Barwood, M., Corbett, J., 2018. Central fatigue theory and endurance exercise: Toward an interoceptive model. Neurosci Biobehav Rev 93, 93–107. https://doi.org/10.1016/j.neubiorev.2018.03.024

Miller, E.K., Cohen, J.D., 2001. An integrative theory of prefrontal cortex function. Annu. Rev. Neurosci. 24, 167–202. https://doi.org/10.1146/annurev.neuro.24.1.167

Miller, M.W., Dacelar, M.F.B., Feiss, R.S., Daou, M., Alderman, B.L., In Press. P3b as an electroencephalographic index of automatic associations of exercise-related images. Int. J. Psychophysiol. http://doi.org/10.1016/j.ijpsycho.2020.10.004

Moreau, D., Chou, E., 2019. The Acute Effect of High-Intensity Exercise on Executive Function: A Meta-Analysis. Perspect. Psychol. Sci. 14, 734–764. https://doi.org/10.1177/1745691619850568

Morres, I.D., Hatzigeorgiadis, A., Stathi, A., Comoutos, N., Arpin-Cribbie, C., Krommidas, C., Theodorakis, Y., 2019. Aerobic exercise for adult patients with major depressive disorder in mental health services: A systematic review and meta-analysis. Depress. Anxiety 36, 39–53. https://doi.org/10.1002/da.22842

O’Leary, K.C., Pontifex, M.B., Scudder, M.R., Brown, M.L., Hillman, C.H., 2011. The effects of single bouts of aerobic exercise, exergaming, and videogame play on cognitive control. Clin Neurophysiol 122, 1518–1525. https://doi.org/10.1016/j.clinph.2011.01.049

Ogoh, S., Ainslie, P.N., 2009. Cerebral blood flow during exercise: mechanisms of regulation. J Appl Physiol (1985) 107, 1370–1380. http://doi.org/10.1152/japplphysiol.00573.2009

Olson, R.L., Chang, Y.K., Brush, C.J., Kwok, A.N., Gordon, V.X., Alderman, B.L., 2016. Neurophysiological and behavioral correlates of cognitive control during low and moderate intensity exercise. NeuroImage 131, 171–180. https://doi.org/10.1016/j.neuroimage.2015.10.011

Polich, J., 2007. Updating P300: an integrative theory of P3a and P3b. Clin Neurophysiol 118, 2128–2148. https://doi.org/10.1016/j.clinph.2007.04.019

Pontifex, M.B., Hillman, C.H., 2007. Neuroelectric and behavioral indices of interference control during acute cycling. Clin Neurophysiol 118, 570–580. https://doi.org/10.1016/j.clinph.2006.09.029

Pontifex, M.B., Parks, A.C., Henning, D.A., Kamijo, K., 2015. Single bouts of exercise selectively sustain attentional processes. Psychophysiology 52, 618–625. https://doi.org/10.1111/psyp.12395

Rogers, P.J., Brunstrom, J.M., 2016. Appetite and energy balancing. Physiol Behav 164, 465–471. https://doi.org/10.1016/j.physbeh.2016.03.038

Schubert, M.M., Desbrow, B., Sabapathy, S., Leveritt, M., 2013. Acute exercise and subsequent energy intake. A meta-analysis. Appetite 63, 92–104. https://doi.org/10.1016/j.appet.2012.12.010

Scudder, M.R., Drollette, E.S., Pontifex, M.B., Hillman, C.H., 2012. Neuroelectric indices of goal maintenance following a single bout of physical activity. Biol. Psychol. 89, 528–531. https://doi.org/10.1016/j.biopsycho.2011.12.009

Selya, A.S., Rose, J.S., Dierker, L.C., Hedeker, D., Mermelstein, R.J., 2012. A practical guide to calculating cohen’s f(2), a measure of local effect size, from PROC MIXED. Front Psychol 3, 111. https://doi.org/10.3389/fpsyg.2012.00111

Sim, A.Y., Wallman, K.E., Fairchild, T.J., Guelfi, K.J., 2014. High-intensity intermittent exercise attenuates ad-libitum energy intake. Int J Obes (Lond) 38, 417–422. https://doi.org/10.1038/ijo.2013.102

Smith, J.L., Carbine, K.A., Larson, M.J., Tucker, L.A., Christensen, W.F., LeCheminant, J.D., Bailey, B.W., 2020. To play or not to play? The relationship between active video game play and electrophysiological indices of food-related inhibitory control in adolescents. https://doi.org/10.31234/osf.io/pby7f

Smith, K.J., Ainslie, P.N., 2017. Regulation of cerebral blood flow and metabolism during exercise. Exp. Physiol. 102, 1356–1371. https://doi.org/10.1113/Ep086249

Spitoni, G.F., Ottaviani, C., Petta, A.M., Zingaretti, P., Aragona, M., Sarnicola, A., Antonucci, G., 2017. Obesity is associated with lack of inhibitory control and impaired heart rate variability reactivity and recovery in response to food stimuli. Int. J. Psychophysiol. 116, 77–84. https://doi.org/10.1016/j.ijpsycho.2017.04.001

Stoet, G., 2010. Sex differences in the processing of flankers. Q. J. Exp. Psychol. (Hove.) 63, 633–638. http://doi.org/10.1080/17470210903464253

Stroth, S., Kubesch, S., Dieterle, K., Ruchsow, M., Heim, R., Kiefer, M., 2009. Physical fitness, but not acute exercise modulates event-related potential indices for executive control in healthy adolescents. Brain research 1269, 114–124. https://doi.org/10.1016/j.brainres.2009.02.073

Swift, D.L., McGee, J.E., Earnest, C.P., Carlisle, E., Nygard, M., Johannsen, N.M., 2018. The Effects of Exercise and Physical Activity on Weight Loss and Maintenance. Prog Cardiovasc Dis 61, 206–213. https://doi.org/10.1016/j.pcad.2018.07.014

Themanson, J.R., Hillman, C.H., 2006. Cardiorespiratory fitness and acute aerobic exercise effects on neuroelectric and behavioral measures of action monitoring. Neuroscience 141, 757–767. https://doi.org/10.1016/j.neuroscience.2006.04.004

Tomporowski, P.D., Ellis, N.R., 1986. Effects of exercise on cognitive processes: A review. Psychol Bull 99, 338–346. https://doi.org/10.1037/0033-2909.99.3.338

Tsai, C.L., Wang, C.H., Pan, C.Y., Chen, F.C., Huang, S.Y., Tseng, Y.T., 2016. The effects of different exercise types on visuospatial attention in the elderly. Psychol. Sport Exerc. 26, 130–138. https://doi.org/10.1016/j.psychsport.2016.06.013

Tsai, C.L., Wang, C.H., Pan, C.Y., Chen, F.C., Huang, T.H., Chou, F.Y., 2014. Executive function and endocrinological responses to acute resistance exercise. Front. Behav. Neurosci. 8. https://doi.org/10.3389/fnbeh.2014.00262

Van Veen, V., Carter, C.S., 2002. The timing of action-monitoring processes in the anterior cingulate cortex. J Cogn Neurosci 14, 593–602. https://doi.org/10.1162/08989290260045837

Watson, T.D., Garvey, K.T., 2013. Neurocognitive correlates of processing food-related stimuli in a go/no-go paradigm. Appetite 71, 40–47. https://doi.org/10.1016/j.appet.2013.07.007

Wellman, P.J., 2000. Norepinephrine and the control of food intake. Nutrition 16, 837–842. https://doi.org/10.1016/s0899-9007(00)00415-9

Wohlwend, M., Olsen, A., Håberg, A.K., Palmer, H.S., 2017. Exercise intensity-dependent effects on cognitive control function during and after acute treadmill running in young healthy adults. Front. Psychol. 8, 406–406. https://doi.org/10.3389/fpsyg.2017.00406

Xie, L., Ren, M., Cao, B., Li, F., 2017. Distinct brain responses to different inhibitions: Evidence from a modified Flanker Task. Sci. Rep. 7, 6657. https://doi.org/10.1038/s41598-017-04907-y

Yanagisawa, H., Dan, I., Tsuzuki, D., Kato, M., Okamoto, M., Kyutoku, Y., Soya, H., 2010. Acute moderate exercise elicits increased dorsolateral prefrontal activation and improves cognitive performance with Stroop test. NeuroImage 50, 1702–1710. https://doi.org/10.1016/j.neuroimage.2009.12.023

